# Evaluating the role of pre-training dataset size and diversity on single-cell foundation model performance

**DOI:** 10.1101/2024.12.13.628448

**Authors:** Alan DenAdel, Madeline Hughes, Akshaya Thoutam, Anay Gupta, Andrew W. Navia, Nicolo Fusi, Srivatsan Raghavan, Peter S. Winter, Ava P. Amini, Lorin Crawford

## Abstract

The success of transformer-based foundation models on natural language and images has motivated their use in single-cell biology. Single-cell foundation models have been trained on increasingly larger transcriptomic datasets, scaling from initial studies with 1 million cells to newer atlases with over 100 million cells. This study investigates the role of pre-training dataset size and diversity on the performance of single-cell foundation models on both zero-shot and fine-tuned tasks. Using a large corpus of 22.2 million cells, we pre-train a total of 400 models, which we evaluate by conducting 6,400 experiments. Our results show that current methods tend to plateau in performance with pre-training datasets that are only a fraction of the size of current training corpora.

## Introduction

The development of single-cell transcriptomic assays has empowered the dissection of biological heterogeneity at multiple scales. Machine learning models have been effectively applied to a range of tasks using single-cell data as input, including batch integration [1, 2], imputation [3], perturbation prediction [4, 5], and reference mapping [6]. In parallel, large general-purpose neural network models, known as foundation models, have revolutionized the fields of natural language processing (NLP) and computer vision [7]. They have even led to groundbreaking progress in biological research domains such as protein modeling [8–10]. Foundation models typically follow a transfer learning paradigm where the model is trained on a generic task, often predicting masked tokens, and then fine-tuned for specific tasks of interest [7]. Notably, recent NLP foundation models have shown strong performance in zero-shot and few-shot scenarios, where no task-specific fine-tuning is performed [11].

In single-cell biology, foundation models hold the potential to enable complex analyses without requiring large amounts of new data for each new scientific question. Recently, several foundation models for single-cell transcriptomics have been developed, including scBERT [12], Geneformer [13], scGPT [14], and scFoundation [15]. Although these models differ in their tokenization and training schemes, most rely on the transformer architecture with self-attention [16]. Typically trained on tens of millions of single-cell transcriptomes, these models are applied to a wide-range of tasks, including batch integration across different sequencing technologies [14], cell type annotation in previously unseen experimental datasets [12, 14, 15], inference of gene regulatory networks underlying tissue-specific mechanisms [13], and generation of gene expression profiles to simulate transcriptomic states following exogenous perturbations [13–15]. The ambition of these single-cell foundation models (scFMs) is to achieve state-of-the-art performance across these tasks and beyond [17]. While foundation models in NLP and computer vision exhibit remarkable zero- and few-shot capabilities [11, 18], the extent to which scFMs can demonstrate analogous levels of performance remains uncertain, as they have been shown to underperform relative to simple baselines [19–21].

Pre-training transformer-based models typically requires expensive, specialized hardware such as graphics processing units (GPUs), tensor processing units (TPUs), or neural processing units (NPUs), as well as days to weeks of time for computation. To date, many scFMs are closed-source or only provide source code for fine-tuning and evaluation, rather than releasing the source code needed to pre-train a model from scratch. Keeping these computational and practical challenges in mind, studies in NLP have demonstrated that more data is not always better [22]. For a fixed training budget, curating a high-quality subset of the data and making additional training passes on that subset may yield better model performance [22]. Although there is some evidence suggesting that current large language models may be under-trained [23, 24], this phenomenon has not yet been evaluated in current scFMs. Nevertheless, developers of scFMs continue to scale up both model and training dataset size. Curating datasets with increased quality and diversity, rather than focusing solely on size, could plausibly improve the performance of scFMs.

In this work, we investigate the role of pre-training dataset size and diversity on the performance of scFMs (Fig. 1). We pre-train a total of 400 models and evaluate them by conducting 6,400 experiments across a comprehensive suite of factors, including both in zero-shot scenarios and when fine-tuned for specific tasks. While many current studies on scFMs focus on assessing their abilities on downstream analyses such as gene network inference [13, 14], model evaluations often lack ground truth labels, making reliable comparisons difficult. Here, we focus on relatively simple, but well-defined, tasks that utilize human-curated labels for classification or aim to predict gene expression directly—namely, batch integration, cell type annotation, and perturbation prediction. Our results indicate that current scFMs do not benefit from increasing pre-training dataset size or from naive approaches to increasing pre-training dataset diversity.

**Figure 1.**
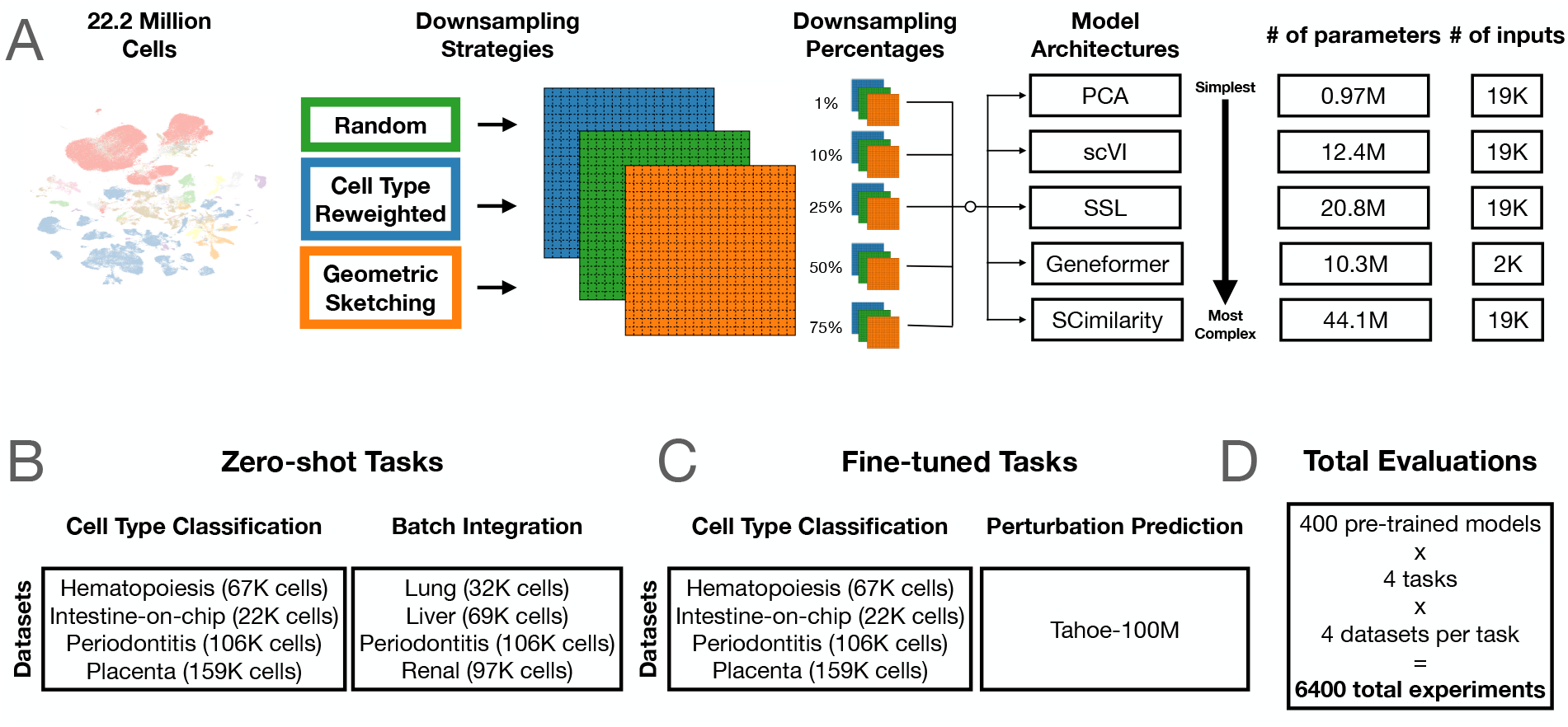
Strategy to assess the effects of pre-training dataset size and diversity on scFM performance. **(A)** Schematic of the downsampling approaches, sizes of downsampled pre-training datasets, and model architectures evaluated. **(B)** Overview of zero-shot tasks and the datasets used for model evaluation. The zero-shot tasks include (i) cell type classification by performing nearest neighbor classification using model embeddings and (ii) an evaluation of how well model embeddings integrate out batch effects. **(C)** Overview of fine-tuned tasks and the datasets used for model evaluation. The finetuned tasks include (i) cell type classification by applying a trained classifier to the model embeddings and (ii) perturbation prediction where a trained regression model is applied to the model embeddings. **(D)** The 400 different pre-trained models were evaluated on 4 different tasks, with each task consisting of 4 different datasets, yielding a total of 6,400 experiments conducted in our main analysis.

## Results

### Overview of pre-training datasets and experimental design

We evaluated the performance of five model architectures of increasing complexity: (1) projection matrices from principal component analysis (PCA) “pre-trained” on a given single-cell dataset, (2) a variational autoencoder (scVI [1]), (3) a masked autoencoder (SSL [25]), (4) a transformer (Geneformer [13]), and (5) an autoencoder-based metric learning approach (SCimilarity [26]). Detailed descriptions of the specific parameters and training procedures for each model architecture are described in the Methods.

We pre-trained each model architecture on datasets of varying size and diversity (Fig. 1A). These pre-training datasets were sampled from the 22.2 million cell scTab corpus of single-cell transcriptomes [27]. To generate these datasets, we implemented three distinct downsampling schemes: (1) random downsampling, which selects cells uniformly at random; (2) cell type re-weighting, which attempts to include equal proportions of each cell type in the dataset; and (3) geometric sketching, which samples cells uniformly across transcriptional space without relying on metadata [28]. Random downsampling conserves diversity (i.e., all subsets have diversity that is approximately equal to the full dataset both in terms of metadata and transcriptomics), while both cell type re-weighting and geometric sketching increase diversity relative to the full corpus. Namely, cell type re-weighting increases diversity with respect to metadata labels, while geometric sketching increases transcriptomic-specific diversity. Pretraining datasets were generated using each of these three downsampling schemes at sizes of 1%, 10%, 25%, 50%, and 75% of the total training corpus. Each pre-training corpus was split into train, validation, and test sets in proportions of 80%, 10%, and 10%, respectively. To ensure robust evaluation, we generated five independent splits for each combination of downsampling scheme and dataset size, resulting in 80 pre-training datasets. All five model architectures were pre-trained on each of these datasets, yielding a total of 80 *×* 5 = 400 pre-trained models for this initial analysis.

### Downsampling schemes increase diversity in pre-training datasets

To confirm that each downsampling scheme had the intended effect on dataset diversity, we computed three diversity metrics for each pre-training dataset: (1) the Shannon index [29], (2) the Gini-Simpson index [30], and (3) the Vendi Score [31]. The Shannon index and the Gini-Simpson index evaluate diversity based on metadata (in this case, the cell type labels), while the Vendi Score directly evaluates the transcriptional diversity. Both cell type re-weighting and geometric sketching increased diversity compared to random downsampling (Supplementary Fig. 1). Interestingly, geometric sketching resulted in higher Vendi Scores compared to cell type re-weighting, whereas cell type re-weighting resulted in higher Shannon and Gini-Simpson index values. These differences reflect the underlying mechanisms of the two downsampling methods: cell type re-weighting enhances diversity based on cell type metadata, while geometric sketching directly targets transcriptional data. The sizes for each dataset range from less than 1 gigabyte (GB) for the 1% datasets to approximately 77 GB for the 100% datasets (Supplementary Fig. 2).

To test whether increased dataset size and/or diversity improved performance (Fig. 1B), we evaluated all 400 pre-trained models in the two primary regimes in which a scFM might be utilized: (1) zero-shot and (2) fine-tuned. In the zero-shot regime, the model’s internal representation of data, referred to as an embedding, is used directly without additional training. In contrast, the fine-tuning regime involves re-training all or part of a model to address a specific task. Fine-tuning typically tackles a supervised learning problem where both labels and the task are well-defined, while the zero-shot approach is suited for tasks where no ground truth labels are available.

### Cell type classification performance plateaus with limited pre-training data

A common task in many single-cell RNA-sequencing (scRNA-seq) workflows is labeling cell types based on their transcriptional profiles. To evaluate initial model performance, we used all pre-trained models to classify cells from a clonal hematopoiesis dataset [32] in both the zero-shot scenario and after fine-tuning the models to classify cell types present in the dataset. For the zero-shot cell type classification task, model embeddings were used directly with a nearest neighbor classifier to classify cells. For the fine-tuned cell type classification task, a neural network classifier was trained using the model embeddings as input.

As simple zero-shot baselines for comparison, we included embeddings derived from using highly variable genes (HVGs) directly and from the top 50 principal components. For both of these embeddings, a nearest neighbor algorithm was used to perform classification, similar to the pre-trained models. For a fine-tuned baseline, we included a regularized logistic classifier trained using HVGs as input features. In both cases, we also implemented “non pre-trained” versions of each model using 0% of the pre-training data. To do this, we simply performed evaluations using each model’s randomly initialized weights. Model performances, as assessed by the micro F1 score, are shown in Fig. 2. Interestingly, in the zeroshot case, Pre-trained PCA, Geneformer, and SCimilarity saw no increase in performance as a function of pre-training dataset size across all downsampling schemes (Fig. 2A). scVI and SSL saw minimal improvement from increasing pre-training dataset size beyond 1%-10% of the entire corpus, except in the case of geometric sketching.

**Figure 2.**
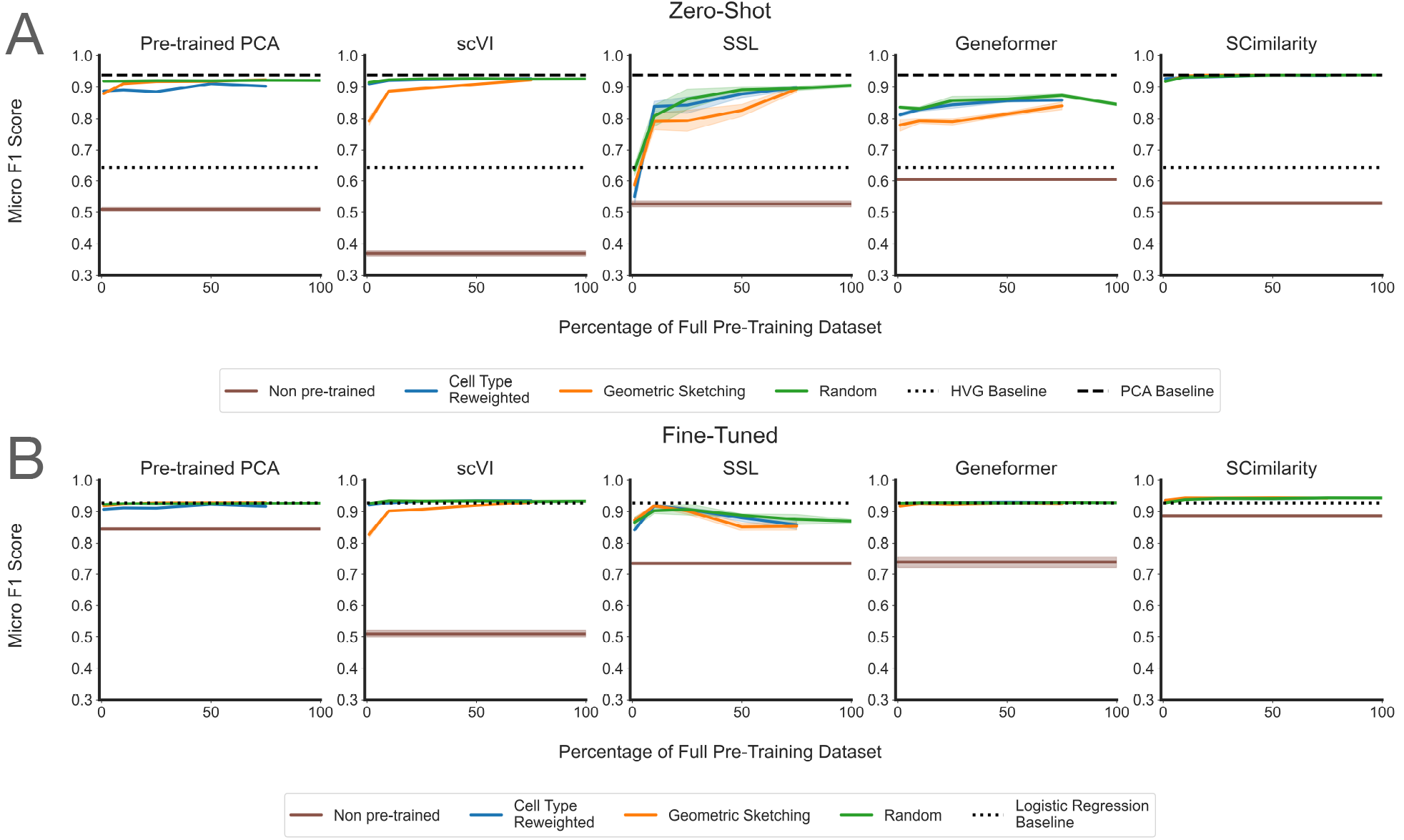
Zero-shot and fine-tuned performance on classifying cells from a clonal hematopoiesis dataset plateaus at a small fraction of the total data available for pre-training. **(A)** Line plots showing zero-shot classification performance for each model’s embeddings, as evaluated by the micro F1 Score. For each model, the different colors correspond to the downsampling strategy used to generate the data used for pre-training. The dotted line shows the performance of using the highly variable genes as an embedding; the dashed line shows the performance of using principal component projections as an embedding; and the brown solid line corresponds to the “non pre-trained” version of each model, where evaluations were done using the randomly initialized weights. **(B)** Line plots showing classification performance for each model after fine-tuning, as evaluated by the micro F1 Score. For each model, the different colors correspond to the downsampling strategy used to generate the data used for pre-training. The dotted line shows the performance of training a regularized logistic regression classifier using the highly variable genes as input features, and the brown solid line corresponds to the “non pre-trained” version of each model, where evaluations were done using the randomly initialized weights. In the fine-tuned tasks, this means that a randomly initialized model is frozen and fine-tuned with no pre-training (as opposed to having no training at all).

When the models had been fine-tuned on the clonal hematopoiesis dataset, this pattern was amplified (Fig. 2B). None of the Pre-trained PCA, SSL, Geneformer, and SCimilarity showed any performance increase from the pre-training dataset size, while scVI exhibited a slight improvement in performance going from 1% of the full corpus to 10% of the full corpus, but no improvement beyond that (except for geometric sketching, which demonstrated reduced performance overall). Notably, we observed a trend similar to recent works [19, 20], where the absolute classification accuracy of more complex methods like SSL and Geneformer was exceeded by both simpler approaches (i.e., Pre-trained PCA and scVI) and baselines (i.e., PCA and logistic regression). While SCimilarity, regardless of downsampling strategy, outperformed all approaches, including the logistic regression baseline, it still did not exhibit improvements from increasing pre-training dataset size. Model performance was comprehensively assessed using accuracy, precision, recall, micro F1 score, and macro F1 score in both the zero-shot scenario (Supplementary Fig. 3) and the fine-tuning scenario (Supplementary Fig. 4). Across all metrics, the phenomenon of model performance plateauing at a small fraction of the total training data was consistently observed. Furthermore, increasing the diversity of pre-training datasets did not improve model performance in this classification task.

### Saturation for the cell type classification task is observed across datasets

To determine whether the performance plateau observed on the clonal hematopoiesis dataset generalized to other datasets, we evaluated cell type classification in both the zero-shot and fine-tuned scenarios across several distinct datasets representing diverse biological contexts: (1) clonal hematopoiesis (as shown in Fig. 2) [32], (2) intestine-on-a-chip [33], (3) placental infection (which we refer to as “placenta”) [34], and (4) periodontitis [35]. Importantly, these datasets are not present in the scTab subset of CELLxGENE and thus represent independent test sets.

For each evaluation dataset, we sought to identify the pre-training corpus fraction where performance gains became marginal, which we refer to as the “learning saturation point” (Fig. 3A). For each combination of model, evaluation dataset, and downsampling scheme, we identified the maximum micro F1 score across all pre-training dataset sizes and defined the “saturation threshold” as 95% of this value. We then identified the smallest percentage of the training data that produced a model surpassing this threshold and labeled this percentage the “learning saturation point”.

**Figure 3.**
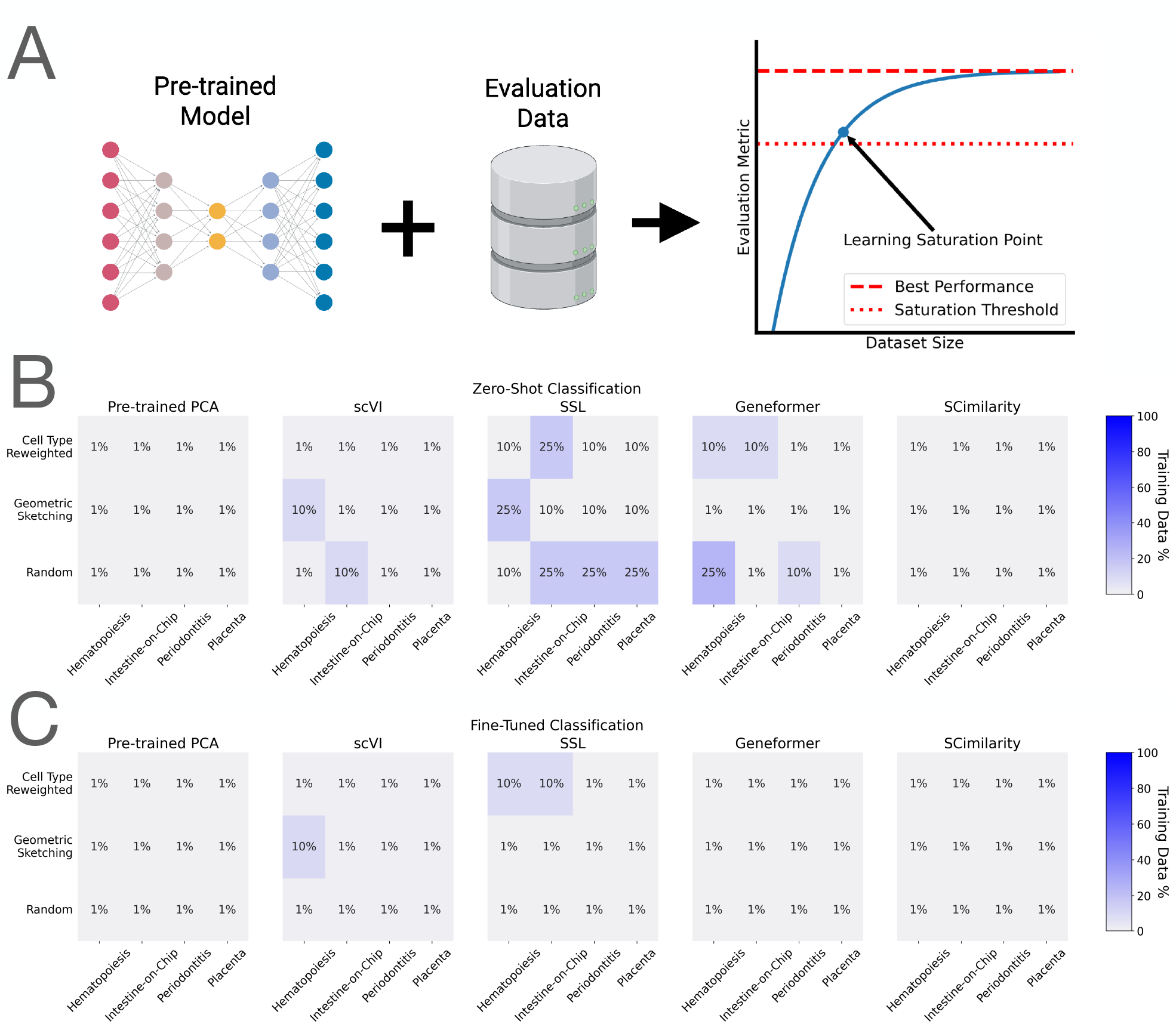
Performance saturation as a function of pre-training dataset size across multiple evaluation datasets on zero-shot and fine-tuned cell type classification. **(A)** Schematic of analysis to find the learning saturation point. For each family of models (i.e., a downsampling strategy paired with a model), a “saturation threshold” of 95% of the maximum performance was computed, and the minimum pre-training dataset size that produced a model surpassing that threshold was identified. This dataset size was denoted the “learning saturation point” and is considered the point at which model performance saturated as a function of pre-training dataset size. **(B)** Heatmap visualizing the learning saturation point for the clonal hematopoiesis, intestine-on-chip, periodontitis, and placental infection datasets for each of Pre-trained PCA, scVI, SSL, Geneformer, and SCimilarity across each downsampling strategy and when evaluated in the zero-shot regime. Each sub-panel corresponds to the model architecture, the *x*-axis corresponds to the dataset evaluated, and the *y*-axis corresponds to the downsampling strategy used to pre-train each model. **(C)** Heatmap visualizing the learning saturation point for the clonal hematopoiesis, intestine-on-chip, periodontitis, and placental infection datasets for each of Pre-trained PCA, scVI, SSL, Geneformer, and SCimilarity, across each downsampling strategy and when evaluated in the fine-tuning regime. Each sub-panel corresponds to the model architecture, the *x*-axis corresponds to the dataset evaluated, and the *y*-axis corresponds to the downsampling strategy used to pre-train each model.

Across all model architectures, downsampling strategies, and evaluation datasets, zero-shot classification performance typically saturated at a small fraction of the total pre-training corpus (Fig. 3B and Supplementary Figs. 5-7), and fine-tuned performance typically saturated at 1% of the training data and never required more than 10% to reach saturation (Fig. 3C and Supplementary Figs. 8-10). For Pre-trained PCA, scVI, and SCimilarity, zero-shot learning saturation largely occurred at 1% and never exceeded 10%; in contrast, for SSL, saturation occurred at 10% or 25% in the zero-shot case. Geneformer overwhelmingly saturated at 1% or 10%, though it reached 25% in one case. Across all four evaluation datasets, more diverse pre-training datasets did not demonstrate improved performance relative to the randomly downsampled datasets. These results suggest that, for cell type classification, the learning saturation point for the tested scFMs can occur as early as 1% of the scTab pre-training corpus, amounting to approximately 200,000 cells. This trend holds for both transformer-based and non-transformer model architectures, as well as for both random and diversity-based downsampling schemes. Notably, 1% and 10% of the total corpus corresponds to less than 1 GB and 9 GB of raw counts, respectively (Supplementary Fig. 2).

### Saturation behavior replicates for zero-shot batch integration

Next, we assessed the performance of the model embeddings on the task of batch integration (Supplementary Figs. 11 and 12). Since obtaining ground truth scRNA-seq data that has been batch effect corrected is not possible, batch integration is inherently a zero-shot task. For this task, model embeddings were evaluated in terms of how well cells of the same type but from different batches were mixed in the embedding space. We utilized four datasets that contained cells from multiple batches, studies, or assays for this evaluation including: (1) the periodontitis dataset [35], previously used in the cell type classification evaluation; (2) a lung dataset [36]; (3) a renal dataset [37]; and (4) a liver dataset [38].

We adopted the batch integration evaluation methodology from Kedzierska et al. [19] and Luecken et al. [39]. This involved generating cluster assignments based on Louvain clusters obtained from the embedding space and comparing them to the ground truth cell type labels. The quality of these cluster assignments was assessed using normalized mutual information (NMI) and adjusted Rand index (ARI). To evaluate the degree of batch mixing in the embedding space, we calculated the average silhouette width (ASW) with respect to both cell type labels and batch labels. The average BIO (AvgBIO) score was then used to summarize overall performance across these metrics, combining ASW, NMI, and ARI. The AvgBIO results for the lung dataset are presented in Fig. 4A, and all metrics for the periodontitis, lung, liver, and renal datasets are shown in Supplementary Figs. 11-14. As observed with cell type classification, the absolute performances of SSL and Geneformer were often exceeded by both simpler approaches (i.e., Pre-trained PCA and scVI) and baselines (i.e., using highly variable genes as embeddings and PCA). The one exception here was, again, SCimilarity, which showed similar performance to scVI.

**Figure 4.**
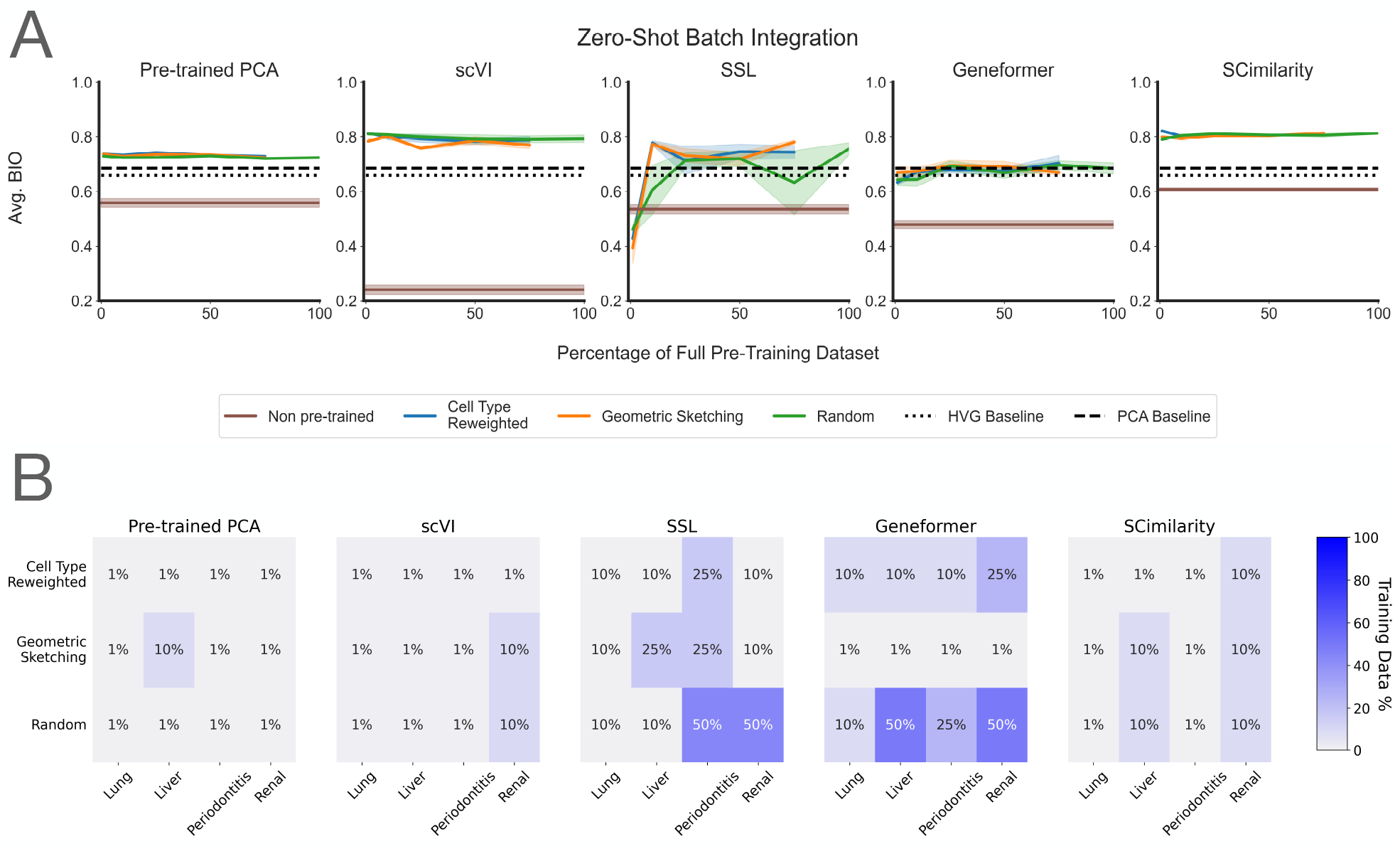
Performance saturation as a function of pre-training dataset size across multiple evaluation datasets on zero-shot batch integration. **(A)** Line plots showing zero-shot batch integration performance for each model’s embeddings, as evaluated by the average BIO (AvgBIO) metric on the lung dataset from Kim et al. [36]. For each model, the different colors correspond to the downsampling strategy used to generate the data used for pre-training. The dotted line shows the performance of using the highly variable genes as an embedding; the dashed line shows the performance of using principal component projections as an embedding; and the brown solid line corresponds to the “non pre-trained” version of each model, where evaluations were done using the randomly initialized weights. **(B)** Heatmap visualizing the learning saturation point for the lung, liver, periodontitis, and renal datasets for each of Pre-trained PCA, scVI, SSL, Geneformer, and SCimilarity, across each downsampling strategy. Each sub-panel corresponds to the model architecture, the *x*-axis corresponds to the dataset evaluated, and the *y*-axis corresponds to the downsampling strategy used to pre-train each model.

We repeated the learning saturation point analysis for the AvgBIO batch integration performance curves across all models, evaluation datasets, and downsampling schemes, and observed that, consistent with the classification task analysis, model performance generally saturated at a small fraction of the total pre-training corpus (Fig. 4B). Specifically, saturation of batch integration performance occurred at 1% for Pre-trained PCA, between 1% and 10% for scVI and SCimilarity, 0% and 50% for SSL, and between 1% and 50% for Geneformer. Notably, in the renal dataset, the “non pre-trained” SSL models performed on par with all of the trained models (Supplementary Fig. 14).

### Saturation behavior replicates for fine-tuned perturbation prediction

Given the saturation behavior observed on both cell type classification and batch integration, we next assessed the performance of the model embeddings on perturbation prediction as a final downstream task (Fig. 5). Although obtaining ground truth scRNA-seq perturbation data for individual cells is not possible, we fine-tuned models on pairs of perturbed and unperturbed cells from the same cell lines (Methods). For this task, we assembled a subset of the large Tahoe-100M experimental atlas [40]. Specifically, we used all cells that were from the (1) LoVo, (2) PANC-1, (3) NCI-H2030, and (4) BT-474 cell lines that were each treated with one of four different drugs: (1) Homoharringtonine, (2) Dinaciclib, (3) PH-797804, (4) TAK-901, or a control treatment of (5) dimethyl sulfoxide (DMSO). To conduct this analysis, we first pre-trained each of PCA, scVI, SSL, Geneformer, and SCimilarity on a downsampled proportion of cells from scTab. During fine-tuning, we then took the model embedding corresponding to an unperturbed cell and trained a simple multi-layer perceptron (MLP) to predict the gene expression of a real perturbed cell of the same type, randomly selected from Tahoe-100M. For each drug, we fine-tuned on three of the four cell lines and left one out for the evaluation. Specifically, we held out LoVo for Homoharringtonine, NCI-H2030 for Dinaciclib, BT-474 for PH-797804, and PANC-1 for TAK-901. Each model was assessed by computing the R-squared (*R*^2^) between the average expression of each individual gene in the actual perturbed cells and the corresponding predicted expression vectors for the same cell type. We also computed the mean squared error (MSE) between the gene expression values in the actual perturbed cells and the predicted expression. The *R*^2^ performance curves for predicting perturbation effects on the LoVo cell line treated with Homoharringtonine are shown in Fig. 5A. The results for all other drugs across the other held out cell lines are similar whether evaluated by either *R*^2^ or mean squared error (Supplementary Figs. 15-18).

**Figure 5.**
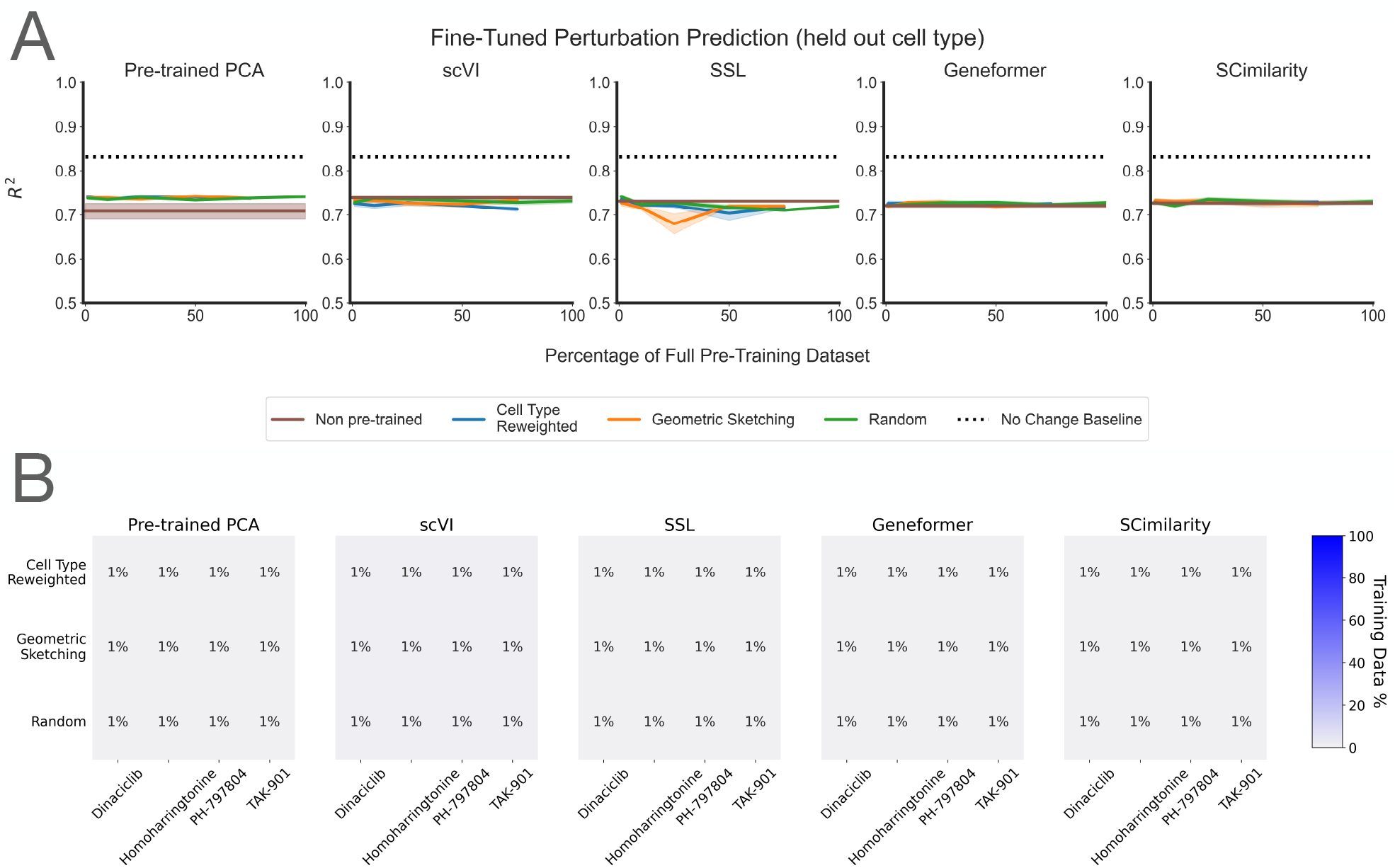
Performance saturation as a function of pre-training dataset size across multiple evaluation datasets on fine-tuned perturbation prediction. **(A)** Line plots showing fine-tuned perturbation prediction performance for each model’s embeddings, as evaluated by *R*^2^ computed between the gene expression for predicted perturbed cells and real perturbed cells. For each model, the different colors correspond to the downsampling strategy used to generate the data used for pre-training. The dotted line shows the performance of using the identity function (i.e., predicting no changed between the unperturbed and perturbed cells); and the brown solid line corresponds to the “non pre-trained” version of each model, where evaluations were done using the randomly initialized weights. **(B)** Heatmap visualizing the learning saturation point for each perturbation dataset, across each downsampling strategy when evaluated on fine-tuned perturbation prediction. Each sub-panel corresponds to the model architecture, the *x*-axis corresponds to the dataset evaluated, and the *y*-axis corresponds to the downsampling strategy used to pre-train each model.

We repeated the learning saturation point analysis for the *R*^2^ performance curves (chosen because it ranges between 0 and 1, in line with the metrics for other tasks) across all models, evaluation datasets, and downsampling schemes. Consistent with the classification task analyses, we observed that model performance generally saturated at a small fraction of the total pre-training corpus (Fig. 5B). Specifically, saturation of perturbation prediction performance occurred at 1% of pre-training dataset size for all model architectures and across all datasets. In line with recent results evaluating deep neural network perturbation prediction performance [21], for all drugs except Dinaciclib (Supplementary Fig. 16), a simple baseline of predicting “no change” in gene expression outperformed each of the fine-tuned models (Supplementary Figs. 15, 17, and 18).

### Assessing the impact of including perturbation data in the pre-training corpus

Since increasing the diversity of the pre-training datasets did not improve model performance on downstream tasks, we reasoned that a small fraction (at maximum, 25%) of the scTab corpus was sufficient to represent cells with a normal transcriptional phenotype. We thus hypothesized that supplementing the pre-training corpus with cells that had been systematically perturbed could enhance the contribution of pre-training on downstream performance.

Here, we performed two sets of spike-in experiments using genome-wide Perturb-seq data from Replogle et al. [41], which contains over 2.5 million systematically perturbed cells. For both sets of experiments, we started with the uniformly downsampled scTab pre-training datasets. For each of these datasets, we generated two new datasets by “spiking in” cells from the Replogle Perturb-seq experiments. This involved replacing a portion of the scTab cells with an equal number of cells from the Replogle Perturb-seq data. The spike-in experiments were conducted at two proportions of spike-in perturbation data: 10% and 50%. The Replogle dataset had enough cells to replace 10% of the scTab cells in all of the uniformly downsampled datasets. However, there were only enough cells to replace 50% of the scTab cells in the 1%, 10%, and 25% downsampled datasets. These experiments allowed us to assess whether incorporating perturbed cells could increase the value of pre-training for downstream tasks. We used these “spike-in” datasets to pre-train scVI, SSL, and Geneformer as representative approaches of a variational autoencoder, a masked autoencoder, and a transformer.

To evaluate the effect of adding in perturbation data, we analyzed the learning saturation points of the micro F1 score performance curves for the cell type classification task, evaluating all models in both the zero-shot (Supplementary Fig. 19) and fine-tuning (Supplementary Fig. 20) cases. All metrics are shown for the zero-shot evaluations in Supplementary Figs. 21-24 and for the fine-tuned evaluations in Supplementary Figs. 25-28. Consistent with the initial analysis of the classification task, model performance generally saturated at a small fraction of the total pre-training corpus. In the fine-tuned case, all model architectures typically saturated at 1%, although Geneformer saturated at 25% for the 10% spike-in in one instance. For scVI, saturation in the zero-shot case typically occurred at 1% and never exceeded 10%. In contrast, SSL saturation occurred between 10% and 25% in the zero-shot case. Finally, Geneformer typically saturated at 1% or 10% in the zero-shot case.

We again repeated the learning saturation point analysis on the AvgBIO performance curves for the batch integration task, evaluating all models pre-trained on the spike-in datasets (Supplementary Fig. 29). All metrics are shown for the zero-shot batch integration evaluations in Supplementary Figs. 30-31. Similar to the classification tasks, across all combinations of model, evaluation dataset, and downsampling schemes, model performance generally saturated at a small fraction of the total pre-training corpus, even with the inclusion of spike-in data. When pre-trained on the spike-in datasets, scVI consistently saturated at 1%, SSL saturated between 10% and 25%, and Geneformer typically saturated at 1% or 10%. From these evaluations in both cell type classification and batch integration, it is clear that naively spiking-in perturbation data does not facilitate an improvement in model performance.

### Despite still saturating with small pre-training datasets, larger models tend to exhibit better overall performance

In each of our previous 6,400 experiments, we held model architectures constant while varying the composition of pre-training datasets. To extend these analyses to the model axis, we explored how model size, hyperparameter configurations, and loss function choices influence downstream predictive performance as the size of a pre-training corpus increases. These evaluations were conducted for zero-shot cell type classification using only the uniformly random downsampled subsets of scTab.

First, to evaluate the role of model size (i.e., the number of parameters in a model), we chose two representative architectures: scVI (representing autoencoders) and Geneformer (representing transformer-based methods). For scVI, we trained models with 0.5*×*, 2*×*, and 4*×* the number of parameters relative to the default implementation by increasing both the size of the two hidden layers and the dimensionality of the latent embedding space. Counting the total number of parameters in Geneformer is more complicated; so, instead, we simply downsized the latent embedding space from the default 256 dimensions to 32, 64, and 128 dimensions. When evaluating scVI, we observed that scaling up the model size resulted in improved downstream performance on the zero-shot classification task across all datasets (Fig. 6A)— albeit, the magnitude of these gains diminished with each successive increase in parameter count. Scaling up Geneformer also improved performance among all evaluation datasets (Fig. 6B), but the architecture with the default 256 latent dimension yielded (approximately) the same accuracy as the model with a latent dimension set to 128. This suggested that a potential plateau in performance could also occur with respect to model size for this particular task. Importantly, while larger models tended to exhibit better overall classification accuracy, none of them benefited from increasing the size of the pre-training dataset.

**Figure 6.**
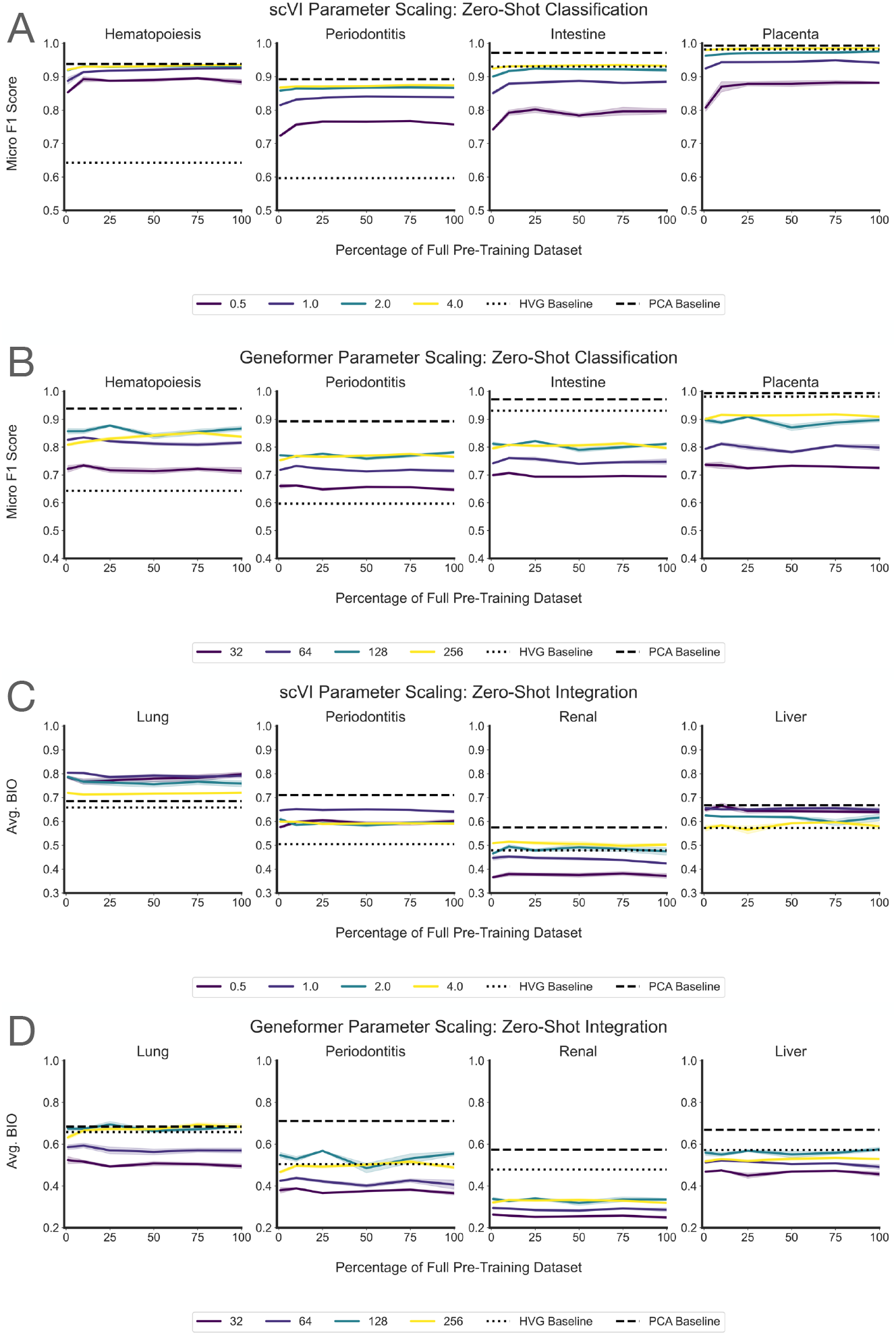
Assessing the impact of model size on zero-shot cell type classification performance. Evaluations were done using the clonal hematopoiesis, intestine-on-chip, periodontitis, and placental infection datasets. **(A)** Line plots showing zero-shot classification performance for scVI models with varying numbers of parameters, as evaluated by the micro F1 Score. For each model, the different colors correspond to the scaling factor for the total number of parameters relative to the default implementation, by scaling both the size of the hidden layers and the dimensionality of the latent embedding space. By default, scVI has a latent embedding space size of 10 and a hidden layer size of 128. In all cases, scVI has one hidden layer in the encoder and one hidden layer in the decoder. **(B)** Line plots showing zero-shot classification performance for Geneformer models with varying numbers of parameters, as evaluated by the micro F1 Score. For each model, the different colors correspond to the dimensionality of the latent embedding space; by default, Geneformer has an embedding size of 256. **(C)** Similar to panel (A) but showing the performance of scVI models with varying numbers of parameters on the zero-shot batch integration task as evaluated by the average BIO (AvgBIO) metric. **(D)** Similar to panel (B) but showing the performance of Geneformer models with varying numbers of parameters on the zero-shot batch integration task as evaluated by the AvgBIO metric.

We also evaluated these models of various sizes on the task of zero-shot batch integration. In contrast to the general result that increasing model scale generally improved zero-shot cell-type classification performance, we observed that for zero-shot batch integration, scVI (Fig. 6C) sometimes showed better performance with a larger model (e.g., the renal dataset), but typically larger models performed worse (e.g., on the lung, periodontitis, and liver datasets). Notably, they were still outperformed by the PCA baseline in the periodontitis, renal, and liver datasets. For Geneformer, the performance on batch integration was quite similar to cell-type classification, where the two larger model sizes showed improved performance; still, however, a latent dimension of 256 did not show benefits over a latent dimension of 128 (Fig. 6D). Further, pre-training dataset size had no impact on model performance across all model scales for both scVI and Geneformer.

For Geneformer, the performance on batch integration was quite similar to cell-type classification and was worse in absolute terms than scVI, reinforcing the observations that Genefomer has poor scaling properties as a function of dataset size and model size (Figs. 2-6). Experiments varying the weight decay hyperparameter had little to no impact on the zero-shot classification (Supplementary Fig. 32A) or zero-shot batch integration (Supplementary Fig. 32B) tasks, underscoring the robustness of these results.

Lastly, since scVI and SSL are both generic autoencoders that use different loss functions, we additionally trained a model we refer to as an “SSL VAE” that integrates the variational loss term of scVI with the masked training objective of SSL. This combined loss function proved more difficult to train, with many models not fully converging in the same number of steps as the original SSL implementation. This resulted in poor performance for zero-shot cell type classification task (Supplementary Fig. 33).

Together, these results demonstrate that, across the autoencoder and transformer architectures, larger models still saturate with small pre-training datasets, though they exhibit better overall performance relative to small models.

## Discussion

In this study, we investigated the role of pre-training dataset size and diversity on the performance of general-purpose single-cell foundation models (scFMs). We evaluated five representative general-use models for single-cell RNA sequencing data on downstream tasks in both the zero-shot case and after finetuning the models for specific tasks. Our work addresses a critical consideration in training large-scale models: the size and diversity of the pre-training corpus. While neural scaling laws observed in other domains suggest that increasing dataset size leads to better model performance, our findings demonstrate that, past a learning saturation point, simply increasing pre-training datasets, even with greater diversity, does not necessarily improve the performance of scFMs on downstream tasks. These results suggest that further scaling up of pre-training datasets from tens of millions of cells to hundreds of millions or even billions of cells may not yield tangible returns.

Importantly, our analysis used the scTab dataset as-is, without additional quality control or data curation beyond downsampling. Recent work has highlighted that some reference atlases contain substantial numbers of low-quality cells and contaminants [42]. Our findings underscore the importance of prioritizing data quality and content over sheer dataset size. This observation is similar to what has been shown in recent work which suggests that performance gains in scFMs can be achieved through principled optimization of smaller training corpora, for example by combining single-cell datasets that capture highly variable, yet unique, transcriptomic states [43]. Our experiments showing that spiking in perturbation data does not necessarily improve performance on cell type classification or batch integration additionally emphasize the importance of using pre-training data and a pre-training task that are aligned with the downstream tasks for which a model will be applied. As such, developers of scFMs and large databases should consider data quality and content rather than simply attempting to gather the largest datasets possible, which we have shown is unlikely to improve performance meaningfully. This is especially important since current foundation models are also quite computationally expensive for both pre-training and inference (Supplementary Fig. 34).

Taken as a whole, our results highlight the need for a more nuanced approach: one that balances dataset size and diversity with careful attention to model architectures, training strategies, and benchmarking. While conceptualizing a cell as “a bag of RNA” [44] is a useful starting point for many models, the concept of a cell as “a sentence of genes” [12–15] may be less effective. Well-designed embeddings have long been used to great success in transcriptomics [1, 45], and simple embeddings can capture a large amount of the information present in single-cell data [19, 46]. Recent work has shown that aligning a model’s training objective with its intended downstream task (e.g., through biologically informed loss functions) can also be an efficient way to enhance its capabilities [47]. In our study, SCimilarity was the best performing model (in absolute terms) at cell type classification, and its pre-training task is heavily aligned with representation learning of cell types [26]. Although transformer-based models have shown great promise, there is a growing body of work [19–21] showing that simple methods often outperform transformer-based models applied to scRNA-seq data. Our results emphasize this fact on multiple tasks and datasets. This suggests that current scFM tokenization schemes that produce “cellular sentences” may be limited in their ability to represent scRNA-seq data faithfully.

Future work should focus on understanding the interplay between feature representation, model architecture, inference, and pre-training dataset size to optimize performance on downstream tasks. Our initial results, which examine the impact of increasing the number of parameters in a model, suggest an interesting path forward. Indeed, large language models have scaling laws that have been established with respect to data, training compute, and model size [48]. Even more, there have been studies focused on developing compute-optimal models that have optimal performance for a given computational budget for training. These frameworks provide guidelines such as “for every doubling of model size the number of training tokens should also be doubled” [49]. The architectures of current scFMs continue to grow in complexity without a complete understanding of where along the compute-data-parameter frontier a given model lies. Although there has been research showing that for scFMs, data quality has a scaling law [50], there has been no comprehensive investigation into computational, data, and parameter size scaling laws for these models. Bridging this gap could unlock principled design strategies for both model architecture development and experimental data collection, as well as accelerate progress toward more generalizable and efficient models for future single-cell applications.

## Methods

### Pre-training data and downsampling schemes

The main pre-training dataset used in this study is scTab [27], which is a subset of the CELLxGENE [51] census (version 2023-05-15). In this particular dataset, there are a total of 22.2 million cells from 164 unique cell types, 5,052 unique human donors, and 56 human tissues. For each of the downsampling strategies described below (uniform random downsampling, cell type re-weighted downsampling, and geometric sketching downsampling), cells in scTab were downsampled in percentages of 1%, 10%, 25%, 50%, and 75% with five replicates for each percentage. Each of these downsampled datasets was then divided into training, validation, and test sets in proportions of 80%, 10%, and 10% splits, respectively. We also downsampled the entire scTab dataset (denoted as the 100% dataset) into 80% training, 10% validation, and 10% test sets with five replicates of this split.

#### Uniform downsampling

Uniform downsampling is the simplest scheme used. The uniformly downsampled datasets sampled each cell uniformly at random without consideration for any additional metadata or the expression counts matrix.

#### Cell type re-weighted downsampling

The goal of the cell type re-weighted scheme is to downsample such that each cell type is present in equal proportions in the resulting dataset. Let *p* = (*p*_1_,…, *p*_*K*_) be the proportion of each cell type in the scTab dataset where *K* is the total number of cell types. For many cell types present in the scTab dataset, there are not enough cells to satisfy this goal even when downsampling to 1%. Therefore, to generate a cell type re-weighted dataset, we first choose all cells that belong to a cell type that is not common enough to satisfy the equal proportion condition. Then, we randomly downsample among the remaining cell types with equal proportions. Let *N* be the total number of cells and *D* be the downsampling proportion. The algorithm is as follows:

1. For any *k*-th cell type satisfying *p*_*k*_*ND < ND/K* (i.e., it is impossible to satisfy the re-weighting condition, even by selecting all of the cells of this cell type), include all of the cells of this cell type.
2. Let *N* ^′^ be the number of unselected cells after the previous step and *K*^′^ be the number of remaining cell types. Some of these cell types that previously satisfied the re-weighting condition may no longer do so with *N* ^′^ and *K*^′^. Repeat step 1 with *N* ^′^ in place of *N* and *K*^′^ in place of *K*.
3. Let *M* be the number of cells already selected. When all remaining cell types satisfy the re-weighting condition, sample (*DN* − *M*)*/K*^′^ cells from each cell type (i.e., re-weight these cell types to have equal proportions).

The resulting downsampled dataset should have proportions 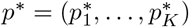 such that 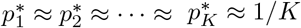.

#### Geometric sketching

Geometric sketching is a method that downsamples based on the intrinsic diversity of a dataset without reference to any metadata [28]. Geometric sketching samples uniformly over transcriptional space without regard to density (i.e., a cell type that has a large number of cells with very similar gene expression will be downweighted in the resulting “sketch”). Let *N* be the total number of cells, *D* be the downsampled proportion, and *C* be the number of covering hypercubes. The geometric sketching algorithm is as follows:

1. Generate a “plaid covering” of the dataset in transcriptomic space. A plaid covering is a covering of the dataset consisting of equally sized hypercubes. Each hypercube in the covering will contain a different number of cells.
2. For each covering hypercube, sample *DN/C* cells.

Due to the fact that each hypercube is of equal size, higher-density regions of transcriptomic space (i.e., regions with many similar cells) will be downweighted in the resulting “sketch” of the dataset.

### Spike-in experiments with perturbation data

In order to assess the extent to which including only normal transcriptomic phenotypes impacts downstream model performance, we performed two sets of “spike-in” experiments using the Replogle et al. [41] Perturb-seq study (CRISPR-based screens with single-cell RNA-sequencing readouts) archived in the scPerturb database [52]. The Replogle dataset contains 2,547,877 cells and 10,117 genes. For both sets of experiments, we began with the downsampled datasets generated using the uniform downsampling scheme. Then, for each of these datasets, we generated two new datasets that had cells from the Replogle Perturb-seq experiments “spiked in”. This was done by removing some of the scTab cells and replacing them with an equal number of cells from the Replogle study. We spiked in these cells at rates of 10% and 50%. The Replogle dataset had enough cells to replace 10% of the scTab cells in all of the uniformly downsampled datasets; however, there were only enough cells to replace 50% of the scTab cells in the 1%, 10%, and 25% downsampled datasets.

### Evaluation datasets

Below is a description of each dataset we used for evaluation in this study. Recall that the clonal hematopoiesis, placenta, intestine-on-chip, and periodontitis datasets were all used for zero-shot and finetuned cell type classification tasks; the lung, renal, and liver datasets were used to evaluate zero-shot batch integration capabilities; and the Tahoe-100M atlas was used to assess perturbation prediction accuracy. All these data, except the lung dataset and Tahoe-100M, were downloaded from the CELLxGENE census (where each of these studies were selected because they published after the 2023-05-15 release and ensured no “leakage” with the training corpora from scTab). The lung dataset was taken from the Curated Cancer Cell Atlas (3CA). Information on where to download these resources can be found in the “Code and data availability” section.

#### Clonal hematopoiesis

The clonal hematopoiesis data results from profiling peripheral blood gene expression from 17 patients with clonal hematopoiesis and 7 controls [32]. For the zero-shot cell type classification evaluations, the data were divided into 50/50 train/test splits. For the fine-tuning evaluations, the data were divided into 80/10/10 train/test/validation splits. There are 66,985 total cells and 8 labeled cell types in this dataset.

#### Placenta

The placenta data results from profiling placental responses to three pathogens [34]. For the zero-shot cell type classification evaluations, the data were divided into 50/50 train/test splits. For the fine-tuning evaluations, the data were divided into 80/10/10 train/test/validation splits. In this dataset, there are 20 donors (counting both mother and fetus), 158,978 total cells, and 7 labeled cell types.

#### Intestine-on-chip

The intestine-on-chip data results from profiling iPSC-derived small intestine-on-chips that were exposed to either: (1) different media conditions or (2) interferon-beta or interferongamma [33]. For the zero-shot cell type classification evaluations, the data were divided into 50/50 train/test splits. For the fine-tuning evaluations, the data were divided into 80/10/10 train/test/validation splits. There are 22,280 total cells and 8 labeled cell types.

#### Periodontitis

The periodontitis data results from profiling the periodontium of 34 individuals [35]. For the zero-shot cell type classification evaluations, the data were divided into 50/50 train/test splits. For the fine-tuning evaluations, the data were divided into 80/10/10 train/test/validation splits. There are 105,918 total cells and 27 labeled cell types.

#### Lung

The lung data results from profiling metastatic lung adenocarcinoma samples from 14 different donors [36]. Note that we do not use training and test splits for the zero-shot batch integration task. Instead, we carry out evaluations using the embeddings from each model directly. There are 32,493 total cells and 10 labeled cell types.

#### Renal

The renal data results from profiling renal cortex samples from 19 different donors [37]. Note that we do not use training and test splits for the zero-shot batch integration task. Instead, we carry out evaluations using the embeddings from each model directly. There are 97,125 total cells and 35 labeled cell types.

#### Liver

The liver data results from profiling the pediatric and adult human liver samples from 16 different donors (7 pediatric, 9 adult) [38]. Note that we do not use training and test splits for the zero-shot batch integration task. Instead, we carry out evaluations using the embeddings from each model directly. There are 69,032 total cells and 19 labeled cell types.

#### Tahoe-100M

The Tahoe-100M is a single-cell atlas of approximately 100 million transcriptomic profiles resulting from 1,100 small molecule perturbations across 50 cancer cell lines [40]. For our analysis, we subset the 1,100 small molecules to consider the four with the greatest median E-distance between perturbed and control cells of the same cellular identity at five micromolar (5 *μ*m), the maximum dosage used in the study (see Figure 4 in Zhang et al. [40] for reference). We then selected four cell lines where at least one of these drugs produced a noticeable non-zero effect (again based on E-distance): PANC-1 for Dinaciclib, BT-474 for Homoharringtonine, NCI-H2030 for PH-797804, and LoVo for TAK-901. For each drug, we fine-tuned using the cell line with the strongest perturbation response and two of the remaining three cell lines, while holding out the fourth for evaluation. Specifically, for Dinaciclib the held out cell line was NCI-H2030, for Homoharringtonine the held out cell line was LoVo, for PH-797804 the held out cell line was BT-474, and for TAK-901 the held out cell line was PANC-1. For perturbation evaluations, the three cell lines used for fine-tuning were divided into 90/10 train/validation splits. In our analyses, control (i.e., unperturbed) cells were cells treated with dimethyl sulfoxide (DMSO). There were far more control cells than treated cells in each dataset, so we additionally subsetted down to 20,000 control cells in the training data and 2,000 cells in the validation data.

### Pre-training of machine learning models

In this study, we implemented five machine learning methods for single-cell transcriptomics: principal component analysis, scVI [1], SSL [25], Geneformer [13], and SCimilarity [26]. For each model class, we standardized the amount of compute (in terms of training steps) that was allocated to pre-training so that simply increasing the pre-training dataset size did not increase the amount of training performed. To determine the number of training epochs allocated to each model, we split the entire scTab corpus into an 80/10/10 split of train, test, and validation datasets, respectively. We then trained each model class on the training split until the validation loss had converged. Next, we plotted the validation loss as a function of the number of epochs and selected the number of epochs that resulted in convergence of the loss function. Finally, this value was normalized based on the fraction of training data used in order to determine the number of epochs needed for each training split. For example, if the number of epochs taken to converge on the full corpus was 40 and we want to find the number of epochs for 25% of the corpus, then we would use 40/0.25 = 160 epochs for this split of the data. In addition to pre-training on corpora generated by each of the downsampling strategies described above (uniform random downsampling, cell type re-weighted downsampling, and geometric sketching downsampling) at percentages of 1%, 10%, 25%, 50%, and 75%, we also implemented “non pre-trained” baseline versions of each model (i.e., using 0% of the pre-training data). To do this, we simply performed evaluations using each model’s randomly initialized weights.

#### Pre-trained principal component analysis (PCA)

To create a “pre-trained” PCA approach, we first subset each pre-training dataset to just include highly variable genes found by using the scanpy package, and then we computed the top 50 principal components of the resulting matrix. For all tasks, each evaluation dataset was projected onto these 50 principal components, and the corresponding vectors were used as embeddings.

#### Single-cell variational inference (scVI)

The default parameters were used to designate the architecture of the pre-trained scVI models. In particular, the number of nodes per hidden layer was 128, the latent embedding space had 10 dimensions, the encoder and decoder both have a single hidden layer, the dropout rate was 10%, the gene likelihood was zero-inflated negative binomial, the dispersion parameter of the negative binomial distribution was constant per gene, and the latent distribution was normal. For training, the batch size was 128, early stopping with respect to the validation loss was turned on, and the number of epochs was determined as 40 *×* (100*/P*) where *P* is the training proportion, since the number of epochs to converge on the full corpus was 40. For example, for a 25 percent split the number of epochs is 40 *×* (100*/*25) = 160 epochs.

#### Self-supervised learning (SSL)

The parameters specified in the SSL repository on GitHub were used to specify the model architecture and pre-training parameters (except for the number of epochs, which was determined as described above). In particular, the hidden layer sizes were (512, 512, 256, 256) for the encoder (reversed for the decoder), and the latent embedding space dimension was 64. For training, the batch size was 16,384, the masking rate was 0.5, the dropout rate was 0.1, the weight decay was set to 0.01, the max steps was 117,000, and the learning rate was 0.001.

#### Geneformer

The parameters specified in the Geneformer repository on Hugging Face were used to specify the Geneformer architecture and pre-training parameters (except for the number of epochs, which was determined as described above with 3 epochs used for the 100% datasets). In particular, the model type was BERT (bidirectional encoder representations from transformers), the maximum input size was 2048, the number of layers was 6, the number of attention heads was 4, the number of embedding dimensions was 256, the intermediate size was 512 (i.e., twice the number of embedding dimensions), and the activation function was ReLU. For training, the batch size was 18 per GPU (across 8 GPUs total), the maximum learning rate was 0.001, the learning schedule was linear, the number of warmup steps was 10,000, the attention dropout was 0.02, the hidden layer dropout was 0.02, the optimizer was AdamW, the layer norm epsilon was 1 *×* 10^−12^, and the initializer range was 0.02, and the weight decay parameter was 0.001.

#### SCimilarity

The parameters specified in the SCimilarity repository on GitHub were used to specify the model architecture and pre-training parameters (except for the number of epochs, which was determined as described above). In particular the hidden layer sizes were (1024, 1024, 1024) for the encoder (reversed for the decoder), the size of the latent dimension was 128, the triplet loss margin was 0.05, the negative selection type was “semihard”, the triplet loss weight was 0.001, the learning rate was 0.005, the batch size was 256, the *T*_max_ for cosine annealing was 0, the dropout was set to 0.5, the input dropout was set to 0.4, the *L*_1_-regularization penalty was 0.0001, and the *L*_2_-regularization penalty was 0.01. The amount of training computation was normalized for all models to a batch size of 1000, 100 batches per epoch, and a maximum of 500 epochs (when pre-training SCimilarity, an epoch is defined not as a pass of the full dataset, but as an arbitrary number of batches used to form triplets).

#### Note on pre-processing datasets for each model

The software for scVI takes in raw gene expression counts. As a result, we did not perform any pre-processing on the pre-training or evaluation data before applying scVI. Similarly, the Geneformer software requires raw gene expression counts before it tokenizes the data; thus, we did not perform any pre-processing before this step in the model. For Pre-trained PCA, SSL, and SCimilarity, gene expression profiles were normalized to 10,000 counts per cell and then logarithmized with a pseudocount of 1 (i.e., processed with the function *f* (**x**) = log(**x** + 1)).

### Experiments assessing the impact of model size

In order to assess the impact of model size on model performance, we performed two sets of model scaling experiments using scVI (because it was the best performing deep generative model in the zeroshot evaluations) and Geneformer (as a representative transformer-based model). To scale scVI, we modulated both the number of nodes per hidden layer and the dimension of the latent embedding space. Note that we kept all other parameters and hyperparameters the same as described above. This generated models that had approximately 0.5*×*, 2*×*, 4*×*, 8*×*, and 16*×* the number of parameters as the default scVI implementation that we highlight in the main text. To scale Geneformer, we modulated the latent embedding space from a default value of 256 to additional values of 32, 64, and 128.

### Experiments assessing the impact of hyperparameter settings

In the next set of experiments, we assessed the impact of different hyperparameter settings on model downstream performance. First, to investigate the influence of weight decay on Geneformer’s downstream performance, we pre-trained different versions of Geneformer, as described above, but with additional weight decay values of 0.01 and 0.0001 (note that the default used was 0.001). Next, we investigated the influence of different training strategies in autoencoder-based models. Both scVI and SSL are autoencoders, where the former uses a variational loss function and the latter makes use of a masked pre-training objective. To assess the relative effects of both approaches, we additionally pre-trained SSL using both the masked training objective and a variational loss term.

### Description of model fine-tuning for cell type classification

Below is a description of how we fine-tune based on the Pre-trained PCA, scVI, SSL, Geneformer, and SCimilarity models.

#### Pre-trained PCA, scVI, and SCimilarity

These three models do not have a fine-tuning mechanism built-in. To get around this fact, we implemented a simple multi-layer perceptron (MLP) that utilized the model embeddings from each of these models as input (this can be conceptualized as freezing the weights of the encoder for each of these models, where PCA is viewed as a linear autoencoder). We utilized an MLP that closely matched the architecture of the scVI decoder, but whose output predicted the cell type label rather than reconstructing gene expression. The model architecture was an input layer of the size of the latent embedding space for the corresponding pre-trained model, a hidden layer of 128 neurons, a ReLU function after the hidden layer, and an output linear layer the size of the number of cell type labels. This architecture was chosen to be simple and was not hyperparameter optimized. For training, we used a dropout rate of 0.2, a batch size of 128, the Adam optimizer with learning rate 1 *×* 10^−4^, cross-entropy loss, early stopping based on the validation loss, and 500 maximum epochs. PCA, scVI, and SCimilarity were trained on heterogeneous hardware consisting of both CPUs and various types of GPUs.

#### Self-supervised learning (SSL)

To fine-tune SSL, we used the built-in cell type classification model from the SSL library. We used the default model architecture where the MLP hidden layers had sizes of 512, 512, 256, 256, and 64. For SSL, no weights were frozen during fine tuning, and for training, we used the AdamW optimizer with a learning rate of 9 *×* 10^−4^, a dropout rate of 0.1, a weight decay of 0.05, a batch size of 16384, a minimum of 30 epochs, used early stopping based on validation loss, and the same number of maximum epochs used during pre-training on a single NVIDIA 80 GB A100 GPU. The learning rate and weight decay were non-default values and were taken from the SSL training scripts.

#### Geneformer

To fine-tune Geneformer, we used the built-in cell type classification model from the Geneformer library. We froze the first 2 layers of the pre-trained model. The fine-tuned MLP classifier adds a final, classification transformer layer. For training, we used a single epoch (following the example set by Theodoris et al. [13]), with 1000 warm-up steps, a linear learning rate schedule, a batch size of 18 per device, and 8 NVIDIA 80 GB A100 GPUs. We used a learning rate 0.000210031 and a weight decay 0.233414 as obtained by using Geneformer’s hyperparameter optimizer.

### Classification tasks and evaluation metrics

We used two simple embeddings as baselines for zero-shot cell type classification: (1) highly variable genes and (2) the cell loadings of the first 50 principal components computed on the dataset. As a baseline for fine-tuned cell type classification, we trained a regularized logistic classifier on the highly variable genes determined on the training dataset. In both cases, for a classification task with cell type labels **y** = (*y*_1_,…, *y*_*N*_) and predicted cell types *ŷ* = (*ŷ* _1_,…, *ŷ*_*N*_), we use the following metrics: accuracy, precision, recall, macro F1 score, and micro F1 score.

#### Accuracy

measures the fraction of cells that are correctly predicted

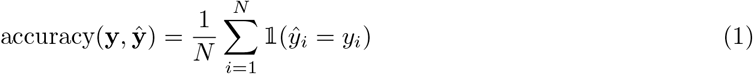

where 𝟙 (•) is an indicator function. For a classification problem, **precision** and **recall** are defined as

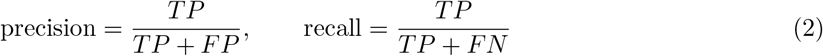

where TP denotes the number of true positives, FP is the number of false positives, and FN is the number of false negatives. Lastly, the **F1 score** is the harmonic mean of the precision and recall such that

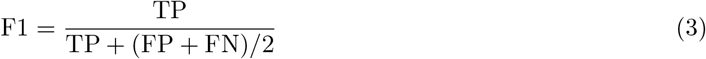

where, in addition to previous notation, TN is the number of true negatives. In this study, we assessed performance in datasets with multiple cell types. There are two variants of the F1 score that measure performance when there are more than two classes. The **micro F1 score** computes TP, FN, and FP globally across all cell types. The **macro F1 score** computes the binary F1 score for each individual label and then averages these F1 scores (which weights each cell type equally, regardless of if they are present in unequal proportions).

### Batch integration task and evaluation metrics

We used two simple embeddings as baselines for zero-shot batch integration: (1) highly variable genes and (2) the cell loadings of the first 50 principal components computed on the dataset. For an integration task with cell type labels **y** = (*y*_1_,…, *y*_*N*_), model embeddings are used to obtain cluster labels **ŷ** = (*ŷ*_1_, …, *ŷ*_*N*_). Here, we use the following metrics: NMI (cluster/label), ARI (cluster/label), ASW (label), ASW (batch), and Average Bio as implemented by Luecken et al. [39] in the `scib`package.

#### Normalized mutual information (NMI)

For two sets of cell type labels, the normalized mutual information (NMI) captures the similarity between the two labels. In the zero-shot integration task, the two labels are the annotated cell type assignments and the clusters derived from the model embeddings. The NMI is computed as follows:

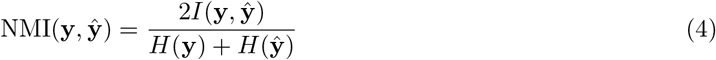

where *I*(**y, ŷ**) is the mutual information of **y** and **ŷ**, and *H*(•) is the Shannon entropy.

#### Adjusted Rand index (ARI)

For two sets of cell type labels, the adjusted Rand index (ARI) captures the similarity between the two labels. In the zero-shot integration task, the two labels are the annotated cell type assignments and the clusters derived from the model embeddings. It is based on the Rand index (RI) which is computed as the following

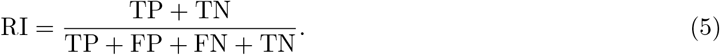

The ARI corrects for the RI measurement’s sensitivity to chance via permutation [53] where

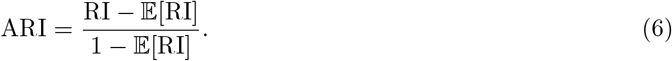

Here, 𝔼 [RI] denotes the expectation of the Rand index.

#### Average silhouette width (ASW)

We compute the average silhouette width considering cell type labels (ASW_*L*_), in order to measure how well embeddings are retaining biological information. We then normalize this score via the following

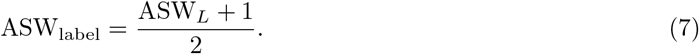

The metric ASW_label_ ranges from 0 to 1 where higher scores indicate better cell type clustering. We also compute the average silhouette width considering batch labels (ASW_*B*_), in order to measure how well embeddings are retaining batch effects. We then invert this score to measure how well embeddings are removing batch effects where

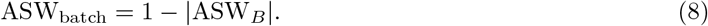

The metric ASW_batch_ also ranges from 0 to 1 where higher scores indicate better batch-mixing performance.

#### Average BIO (AvgBIO)

The AvgBIO score is the arithmetic mean of ASW, NMI, and ARI where

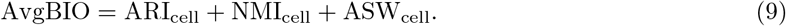

This metric was introduced by Cui et al. [14] and is intended to measure the biological conservation of the embeddings being evaluated. AvgBIO is normalized to the unit scale with 1 indicating complete agreement between clusters and cell type labels.

### Perturbation prediction task and evaluation metrics

To perform fine-tuned perturbation prediction, we implemented the following strategy motivated by Lotfollahi et al. [4]. First, we pre-trained PCA, scVI, SSL, Geneformer, and SCimilarity on some set of unperturbed cells from scTab. During fine-tuning, we took the embeddings corresponding to an unperturbed cell and trained a simple multi-layer perceptron (MLP) to predict the gene expression of a real (and randomly selected) perturbed cell of the same type. The MLP had an architecture with an input layer the size of each model’s embedding dimension (i.e., 50 for Pre-trained PCA, 10 for scVI, 64 for SSL, 256 for Geneformer, and 128 for SCimilarity), a hidden layer of 128 neurons, a ReLU function after the hidden layer, and an output linear layer size equal to the number of cell type labels. Note that this architecture was chosen to be simple and was not optimized. During fine-tuning, we used a dropout rate of 0.2, a batch size of 128, the Adam optimizer with learning rate 1 *×* 10^−4^, cross-entropy loss, early stopping based on the validation loss, and 500 maximum epochs.

When conducting evaluation, we took embeddings corresponding to unseen unperturbed cells from a particular cell line and generated a predicted gene expression vector using the fine-tuned MLP. We then compared these predictions to the average expression of real perturbed cells from the same cell line (on a per-gene basis). We used two metrics to assess perturbation prediction accuracy: (1) **the coefficient of determination** (or R-squared, *R*^2^) and (2) **mean squared error**. All *R*^2^ values were calculated by squaring the *r* output from the `scipy.stats.linregress`function.

### Data diversity measures

We used three different diversity measures to assess the impact of the various downsampling schemes on the diversity of the data used for model pre-training. The Shannon index and Gini-Simpson index are based on cell metadata, while the Vendi Score is based on the gene expression values directly. For the two metadata-based measures, we use the following notation: let *p* = (*p*_1_,…, *p*_*K*_) be the proportion of each cell type in the dataset where *K* is the total number of cell types.

#### Shannon index

The Shannon index is a popular diversity measure in ecology [30] and is the application of Shannon entropy [29] to group proportions. In particular, the Shannon index is given by

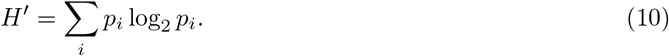

While the Shannon index can be defined using any base for the logarithm, in this study we use base 2. For the Shannon index, a higher value indicates increased diversity.

#### Simpson index and Gini-Simpson index

The Simpson index is also a popular diversity index in ecology [54] and represents the probability that two randomly chosen cells belong to the same cell type

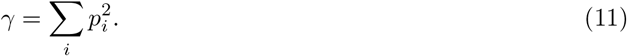

For the Simpson index, a higher value (counter-intuitively) indicates decreased diversity. This leads to the definition of the Gini-Simpson index which is simply the complement of the Simpson index

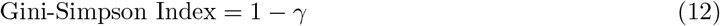

For the Gini-Simpson index, a higher value indicates increased diversity.

#### Vendi Score

The Vendi Score is a measure of the intrinsic diversity of a dataset, without reference to any metadata [31]. In this study, we calculate the Vendi Score of the gene expression matrix for each dataset. When using a linear kernel, the Vendi Score has computational complexity *O*(min(*J*^3^,*N* ^3^)) where *N* is the number of samples in the dataset and *J* is the number of features in the dataset. Due to this, we pre-processed each dataset by

1. Noramalizing cells to 10,000 counts per cell.
2. Logarithmizing the counts with a pseudocount of 1 (i.e., using the function *f* (*x*) = log(*x* + 1)).
3. Restricting to highly variable genes using the `scanpy`function `sc.pp.highly_variable_genes`with default parameters.
4. Centering and scaling the variable genes that remained.

Suppose that we have a collection of cells (**x**_1_,…, **x**_*N*_) and define a positive semidefinite similarity function *k* that satisfies *k*(**x, x**) = 1. Next let **K** be an *N × N* kernel matrix where each entry is *K*_*ij*_ = *k*(**x**_*i*_, **x**_*j*_), and let *λ*_1_,…, *λ*_*N*_ be the eigenvalues of **K***/N* . The Vendi Score with respect to the function *k* is then the exponential of the Shannon entropy of the eigenvalues of **K***/N* where

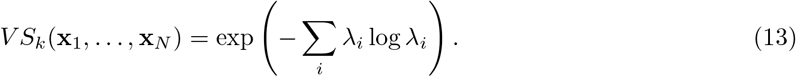

The Vendi Score can also be viewed as the effective rank of a matrix and was previously introduced in the signal processing literature [55].

## Code and data availability

All code is available under an open-source MIT license at https://github.com/microsoft/scFM-dataselection. Specifically, we used scVI [1] which was downloaded from https://scvi-tools.org/ [56] (release 1.1.2), the version of SSL [25] corresponding to commit a149b403fb2f7fe976cb8bd30fcb3d b7a51e9500 (May 2, 2024) from the GitHub repository https://github.com/theislab/ssl_in_scg, the version of Geneformer [13] corresponding to commit b2bbd7ccc856e3b3c1a9199c7cca07d888d87663 (Jun 29, 2024) from the Hugging Face repository https://huggingface.co/ctheodoris/Geneformer/tree/main/geneformer, and SCimilarity (release 0.4.1) from https://github.com/Genentech/scimilarity. The clonal hematopoiesis, placenta, intestine-on-chip, periodontitis, renal, and liver datasets are all available in the CELLxGENE census and can be found online at https://cellxgene.cziscience.com/. The lung dataset from Kim et al. [36] is available in the Curated Cancer Cell Atlas (3CA) and can be found online at https://www.weizmann.ac.il/sites/3CA/lung. The Tahoe-100M dataset is available online via Google Cloud Storage (under bucket gs://arc-ctc-tahoe100/2025-02-25/) and instructions for downloading can be found at https://github.com/ArcInstitute/arc-virtual-cell-atlas/blob/main/tahoe-100M/tutorial-py.ipynb. Instructions for downloading the scTab corpus from the CELLxGENE census are provided at https://github.com/microsoft/scFM-dataselection.

## Acknowledgments

This research was conducted using a combination of computational resources and services provided by Microsoft Research and the Center for Computation and Visualization at Brown University. Parts of Figure 3 were created using BioRender.com. This research was also supported in part by a David & Lucile Packard Fellowship for Science and Engineering awarded to LC. SR acknowledges funding support from NCI K08 CA260442. Any opinions, findings, and conclusions or recommendations expressed in this material are those of the author(s) and do not necessarily reflect the views of any of the funders.

## Author contributions

AD, APA, and LC conceived the study. AD, MH, and AT developed the software and performed the analyses. AG and AWN conducted secondary analyses. NF, SR, PSW, APA, and LC provided resources. SR, PSW, APA, and LC supervised the project. AD, MH, AT, APA, and LC wrote the initial draft. All authors interpreted the results and revised the manuscript.

