## Supplementary Figures for "Evaluating the role of pre-training dataset size and diversity on single-cell foundation model performance"

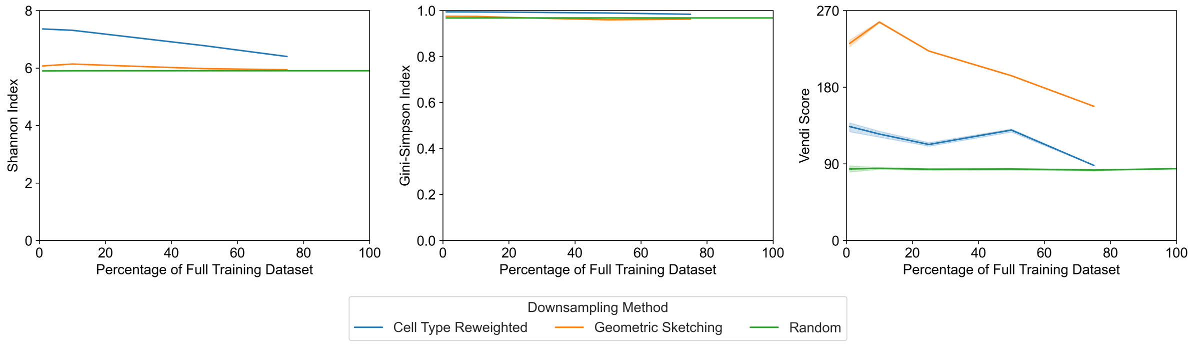


**Supplementary Figure 1. Diversity of datasets used for pre-training as evaluated by intrinsic and extrinsic metrics.** The Shannon index, Gini-Simpson index, and Vendi Score are shown for each of the downsampled pre-training datasets. Cell type re-weighting and geometric sketching have increased diversity relative to the randomly downsampled datasets. Cell type re-weighting (which re-weights based on cell type metadata) has the highest Shannon index and Gini-Simpson index (which both measure the diversity of cell type metadata). Geometric sketching (which samples evenly across transcriptional space) has the highest Vendi Score (which measures the diversity of the transcriptional data directly).

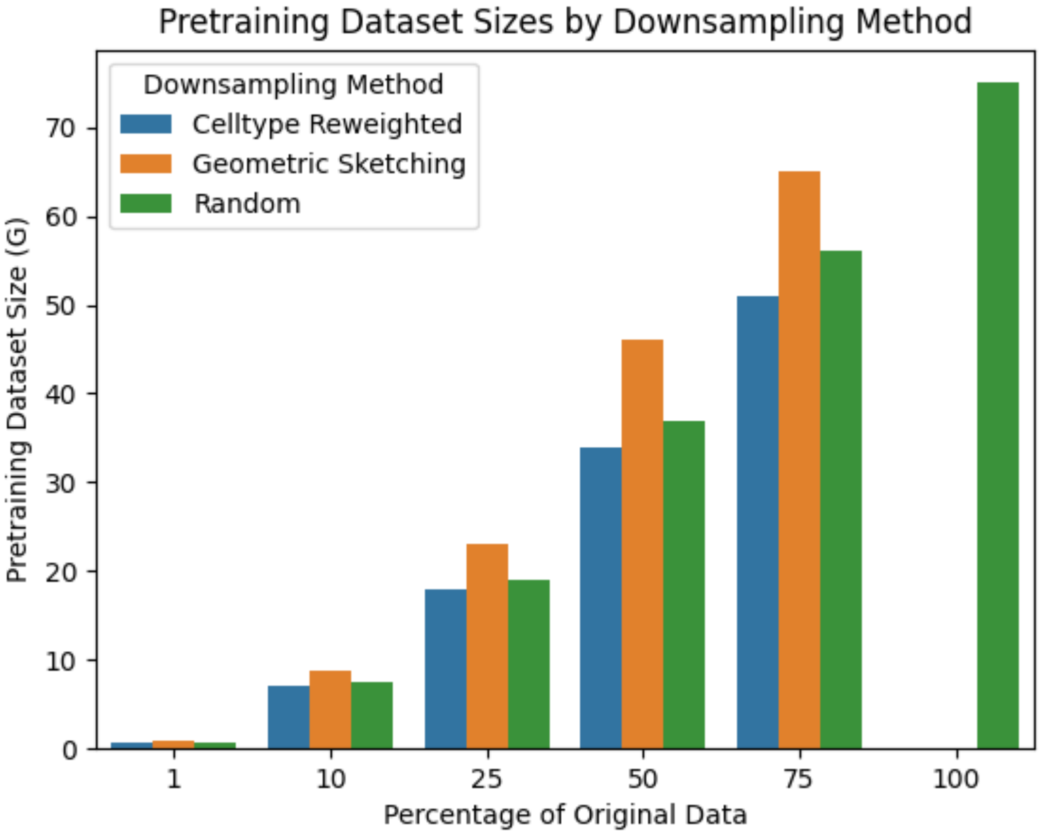


**Supplementary Figure 2. Pre-training dataset sizes by downsampling method.** A barplot showing the dataset size represented via gigabytes (GB) computed using the utility `dh` on the `gzip` compressed `h5ad` files of the single-cell expression matrices.

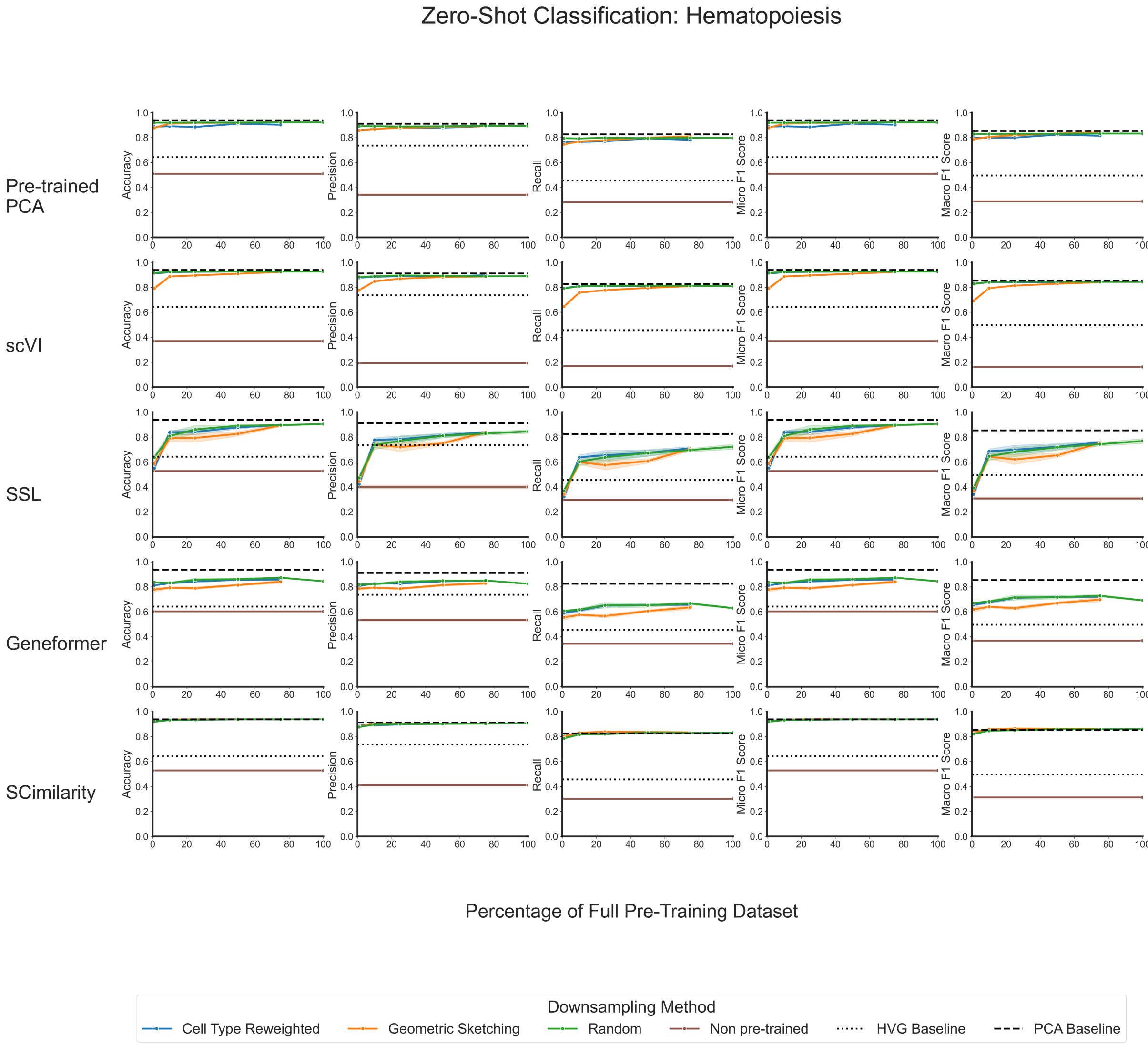


**Supplementary Figure 3. Multiple metrics show that zero-shot model performance on classifying cells from a clonal hematopoiesis dataset plateaus at a small fraction of the total data available for pre-training.** Line plots showing zero-shot classification performance for each model's embeddings as evaluated by accuracy, precision, recall, micro F1 score, and macro F1 score. For each model, the different colors correspond to the downsampling strategy used to generate the data used for pre-training. The dotted line shows the performance of simply using the highly variable genes as an embedding; the dashed line shows the performance of using principal component projections as an embedding; and the brown solid line corresponds to the "non pre-trained" version of each model, where evaluations were done using the randomly initialized weights.

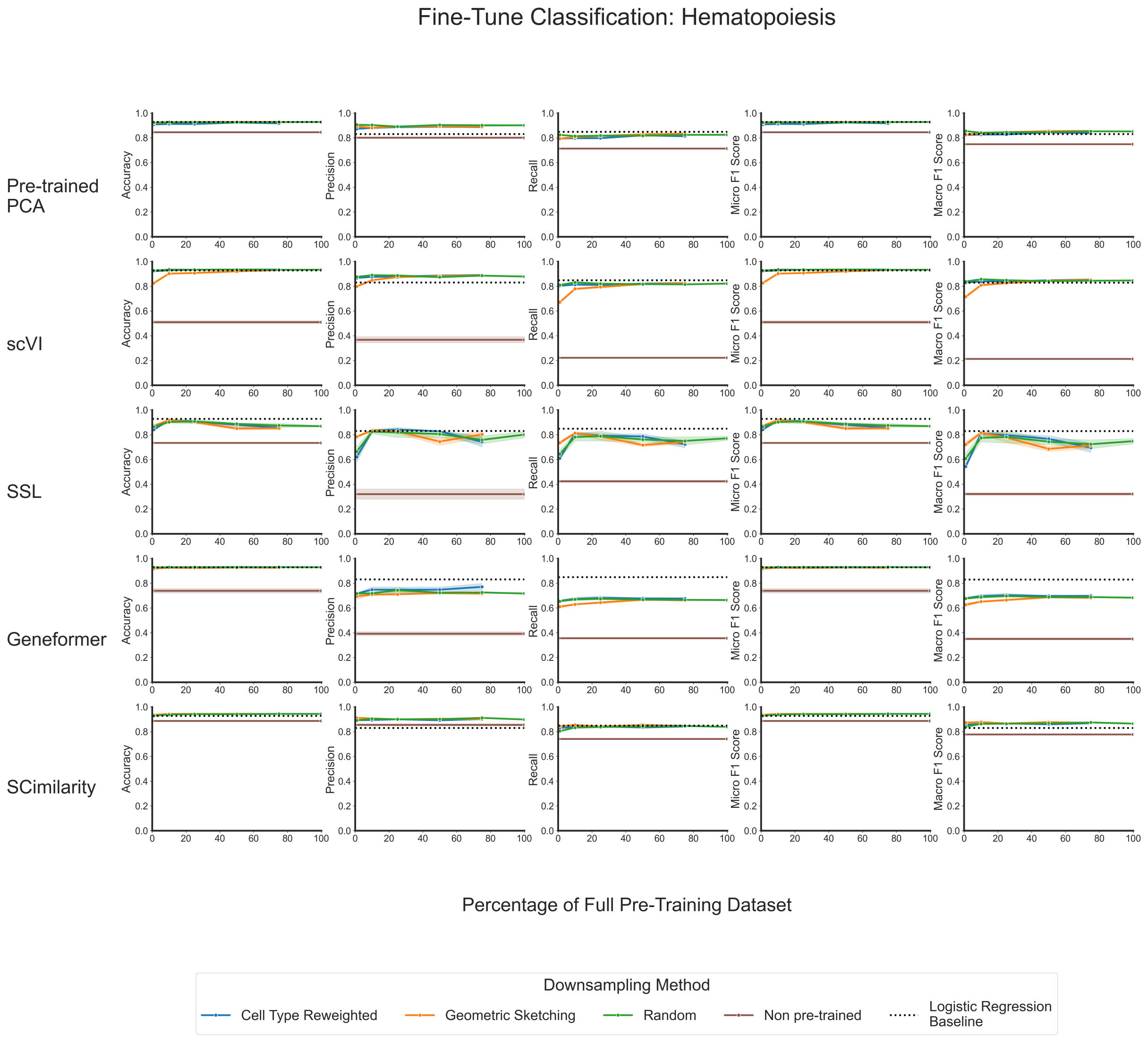


**Supplementary Figure 4. Multiple metrics show that fine-tuned model performance on classifying cells from a clonal hematopoiesis dataset plateaus at a small fraction of the total data available for pre-training.** Line plots showing zero-shot classification performance for each model's embeddings as evaluated by accuracy, precision, recall, micro F1 score, and macro F1 score. For each model, the different colors correspond to the downsampling strategy used to generate the data used for pre-training. The dotted line shows the performance of a regularized logistic classifier using the highly variable genes as input; and the brown solid line corresponds to the "non pre-trained" version of each model, where evaluations were done using the randomly initialized weights.

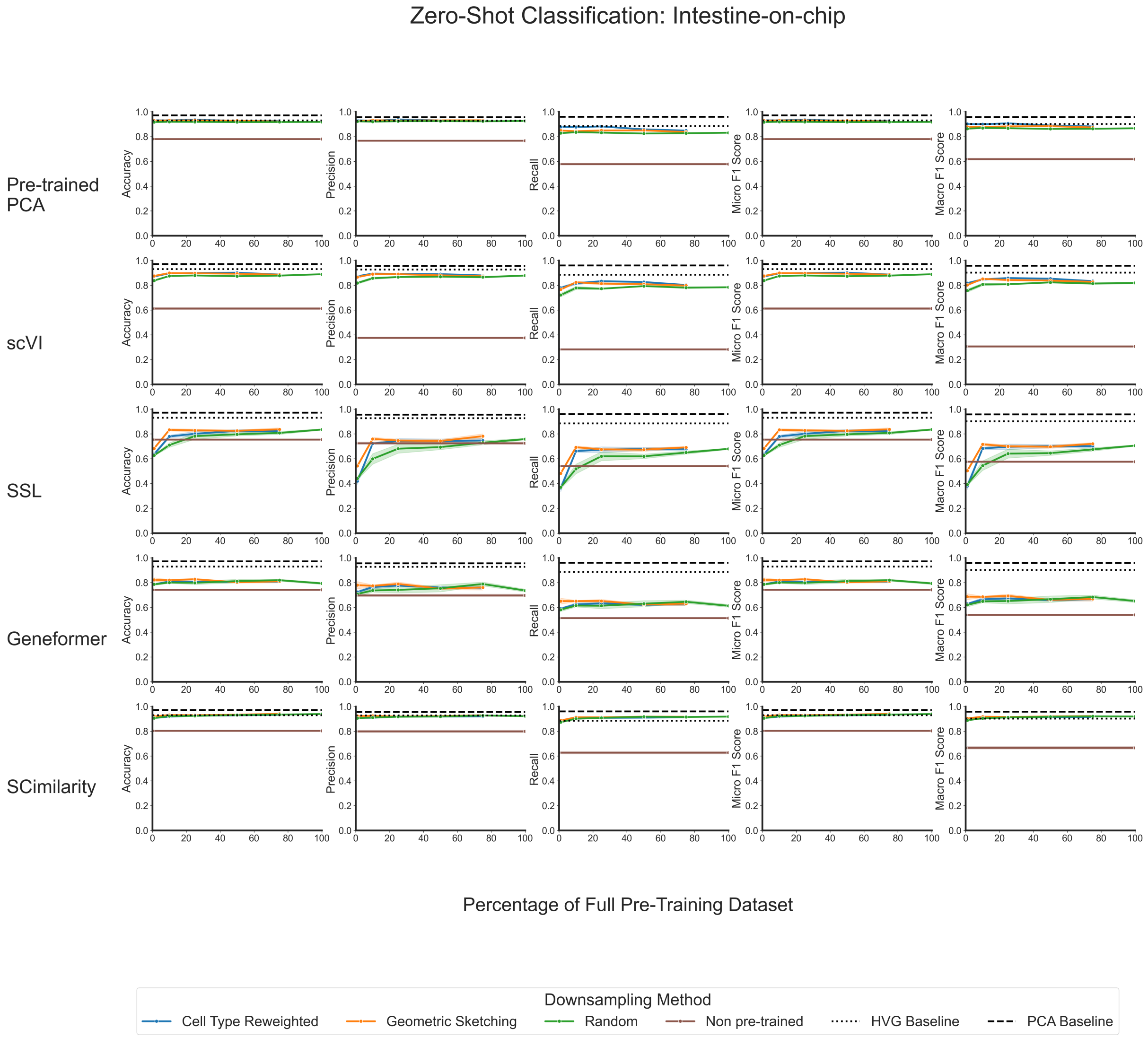


**Supplementary Figure 5. Multiple metrics show that zero-shot model performance on classifying cells from an intestine-on-chip dataset plateaus at a small fraction of the total data available for pre-training.** Line plots showing zero-shot classification performance for each model's embeddings as evaluated by accuracy, precision, recall, micro F1 score, and macro F1 score. For each model, the different colors correspond to the downsampling strategy used to generate the data used for pre-training. The dotted line shows the performance of simply using the highly variable genes as an embedding; the dashed line shows the performance of using principal component projections as an embedding; and the brown solid line corresponds to the "non pre-trained" version of each model, where evaluations were done using the randomly initialized weights.

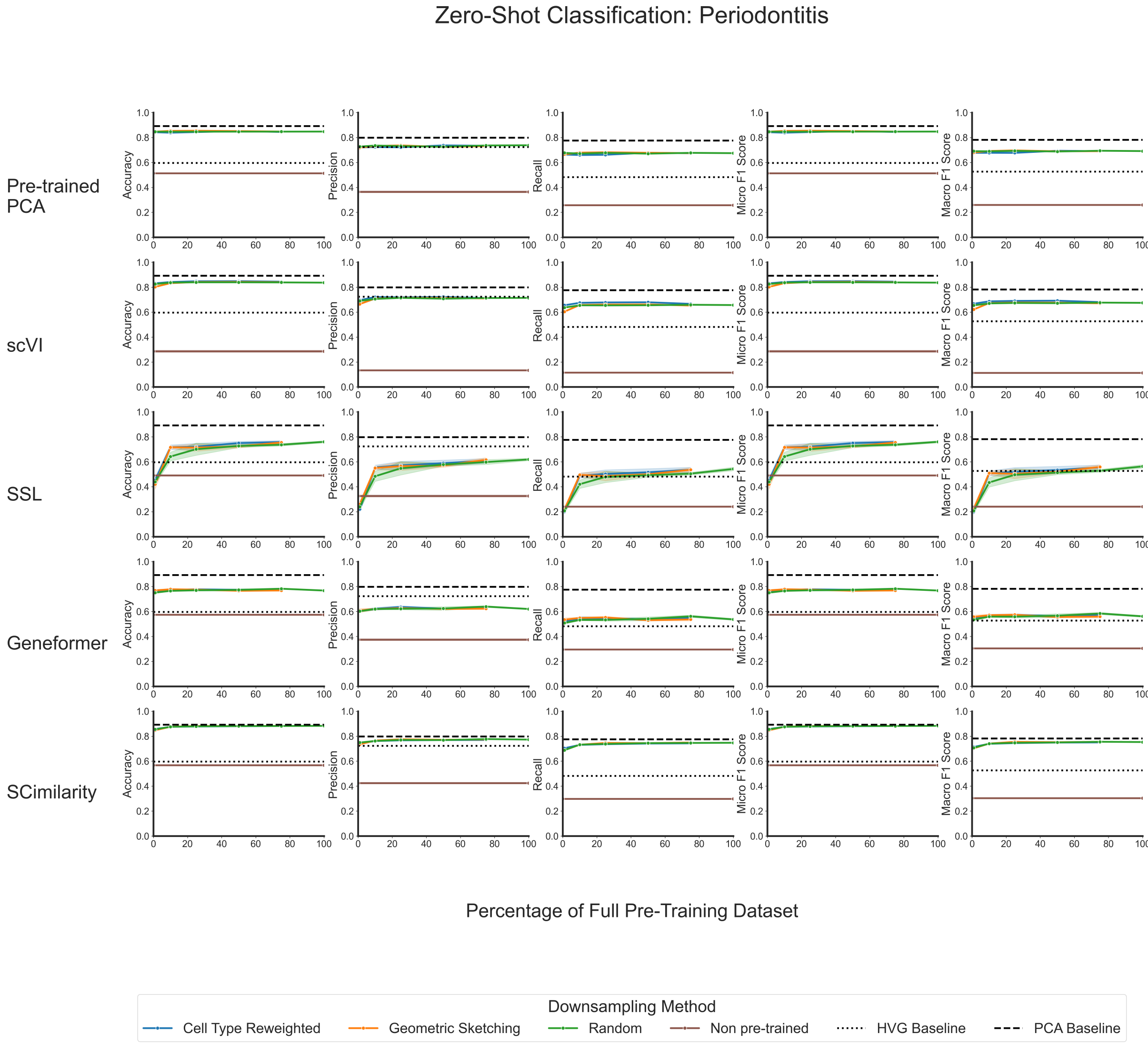


**Supplementary Figure 6. Multiple metrics show that zero-shot model performance on classifying cells from a periodontitis dataset plateaus at a small fraction of the total data available for pre-training.** Line plots showing zero-shot classification performance for each model's embeddings as evaluated by accuracy, precision, recall, micro F1 score, and macro F1 score. For each model, the different colors correspond to the downsampling strategy used to generate the data used for pre-training. The dotted line shows the performance of simply using the highly variable genes as an embedding; the dashed line shows the performance of using principal component projections as an embedding; and the brown solid line corresponds to the "non pre-trained" version of each model, where evaluations were done using the randomly initialized weights.

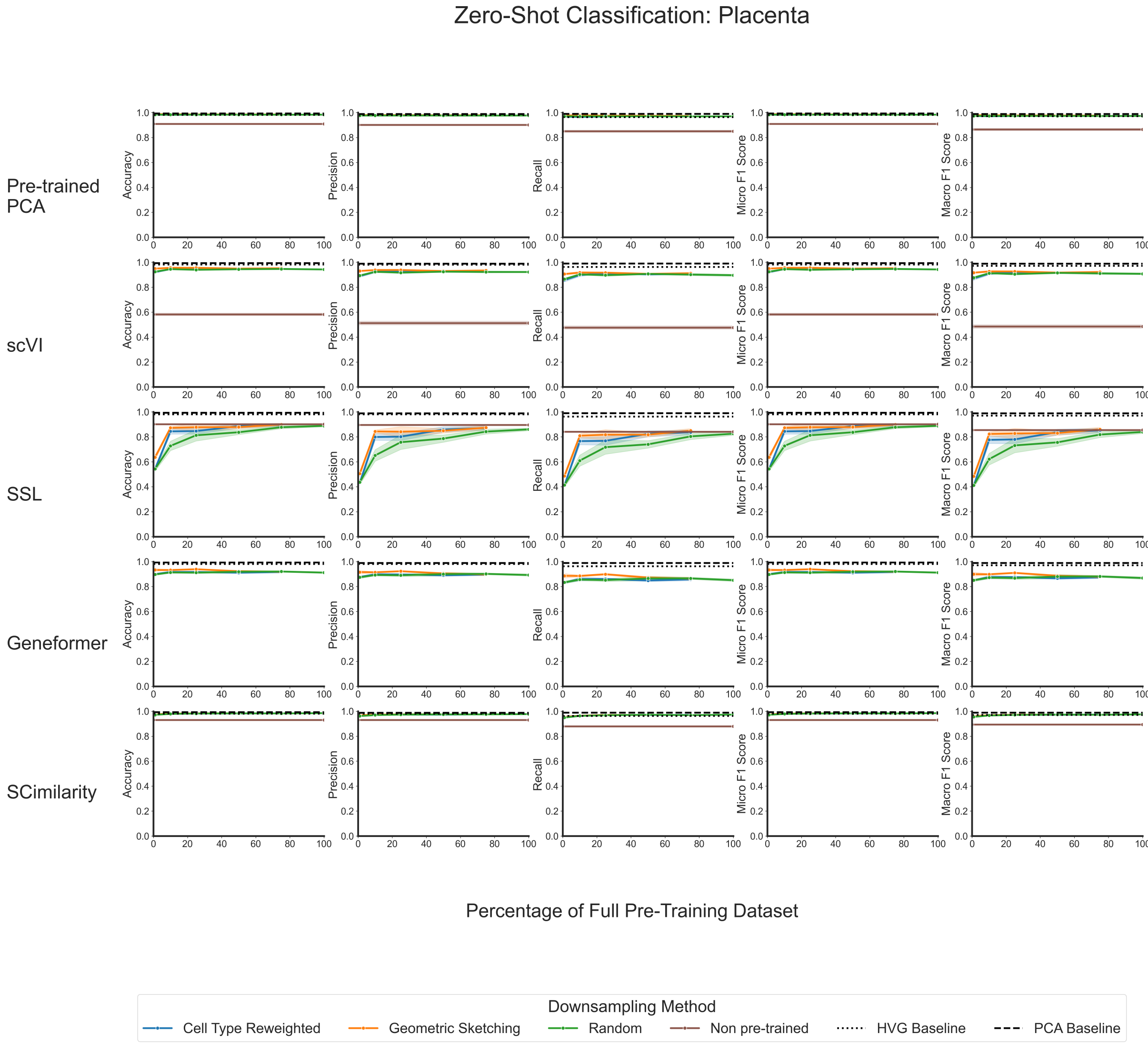


**Supplementary Figure 7. Multiple metrics show that zero-shot model performance on classifying cells from a placenta dataset plateaus at a small fraction of the total data available for pre-training.** Line plots showing zero-shot classification performance for each model's embeddings as evaluated by accuracy, precision, recall, micro F1 score, and macro F1 score. For each model, the different colors correspond to the downsampling strategy used to generate the data used for pre-training. The dotted line shows the performance of simply using the highly variable genes as an embedding; the dashed line shows the performance of using principal component projections as an embedding; and the brown solid line corresponds to the "non pre-trained" version of each model, where evaluations were done using the randomly initialized weights.

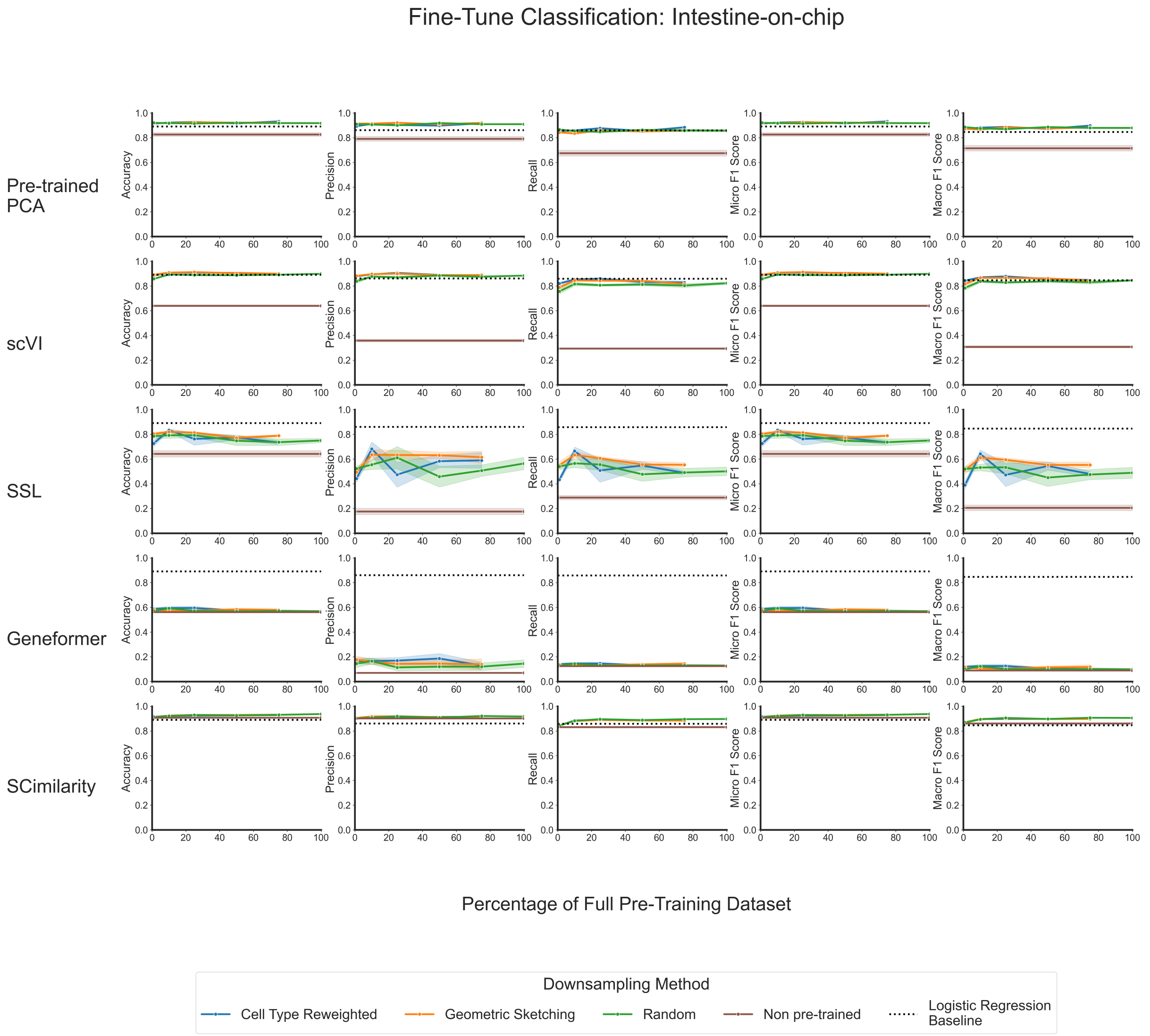


**Supplementary Figure 8. Multiple metrics show that fine-tuned model performance on classifying cells from an intestine-on-chip dataset plateaus at a small fraction of the total data available for pre-training.** Line plots showing zero-shot classification performance for each model's embeddings as evaluated by accuracy, precision, recall, micro F1 score, and macro F1 score. For each model, the different colors correspond to the downsampling strategy used to generate the data used for pre-training. The dotted line shows the performance of a regularized logistic classifier using the highly variable genes as input; and the brown solid line corresponds to the "non pre-trained" version of each model, where evaluations were done using the randomly initialized weights.

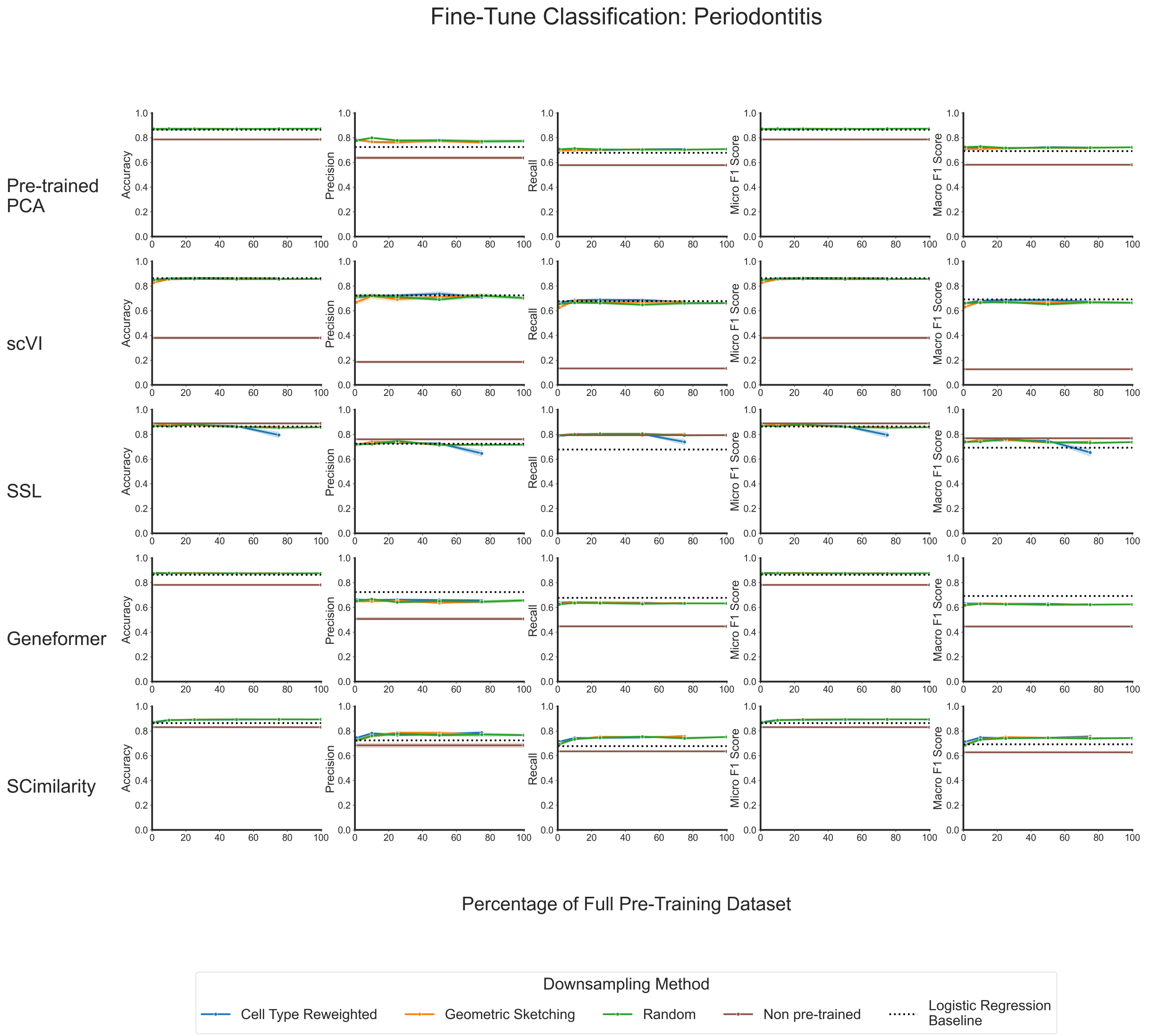


**Supplementary Figure 9. Multiple metrics show that fine-tuned model performance on classifying cells from a periodontitis dataset plateaus at a small fraction of the total data available for pre-training.** Line plots showing zero-shot classification performance for each model's embeddings as evaluated by accuracy, precision, recall, micro F1 score, and macro F1 score. For each model, the different colors correspond to the downsampling strategy used to generate the data used for pre-training. The dotted line shows the performance of a regularized logistic classifier using the highly variable genes as input; and the brown solid line corresponds to the "non pre-trained" version of each model, where evaluations were done using the randomly initialized weights.

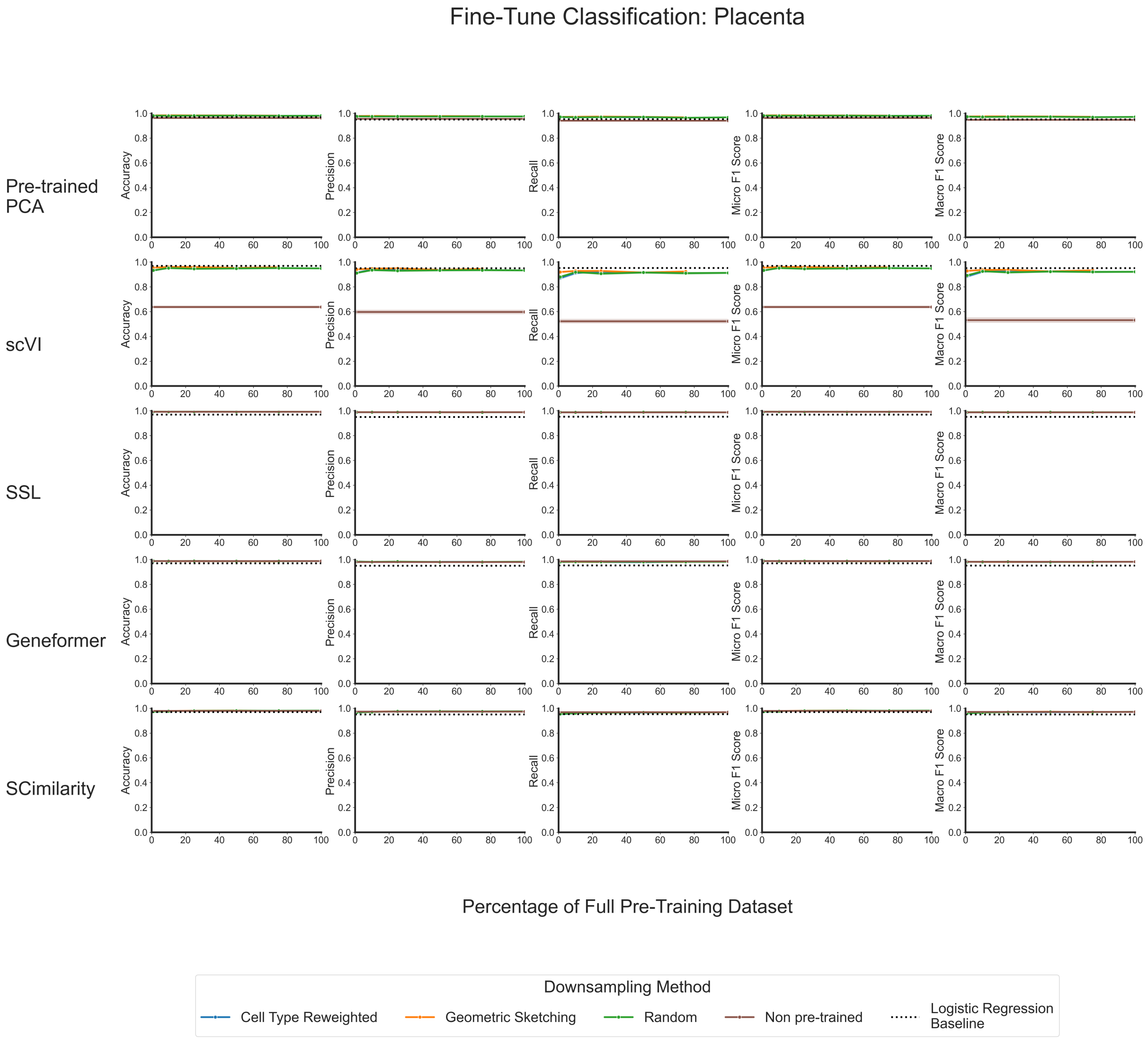


**Supplementary Figure 10. Multiple metrics show that fine-tuned model performance on classifying cells from a placenta dataset plateaus at a small fraction of the total data available for pre-training.** Line plots showing zero-shot classification performance for each model's embeddings as evaluated by accuracy, precision, recall, micro F1 score, and macro F1 score. For each model, the different colors correspond to the downsampling strategy used to generate the data used for pre-training. The dotted line shows the performance of a regularized logistic classifier using the highly variable genes as input; and the brown solid line corresponds to the "non pre-trained" version of each model, where evaluations were done using the randomly initialized weights.

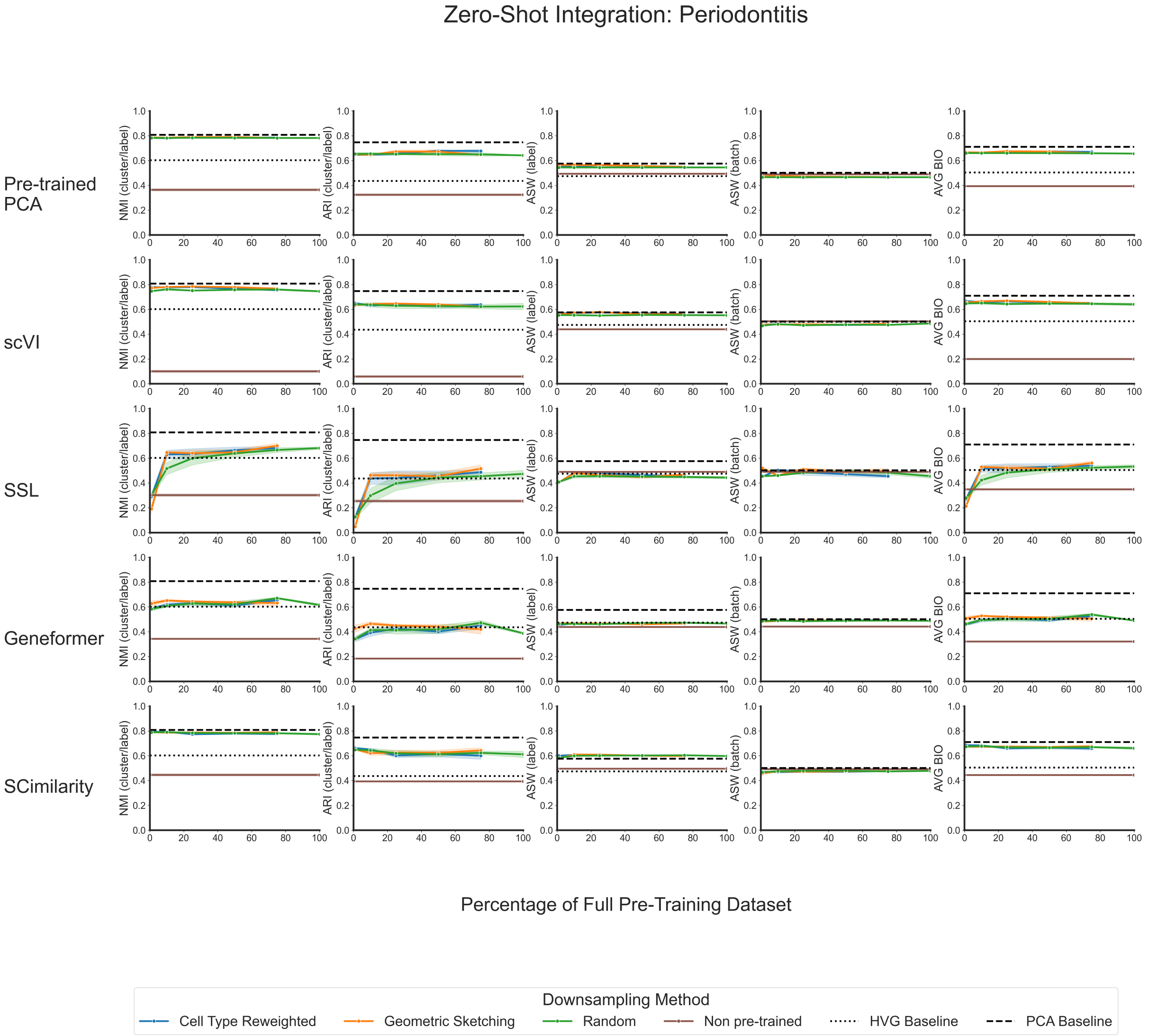


**Supplementary Figure 11. Multiple metrics show that zero-shot model performance on integrating cells from a periodontitis dataset plateaus at a small fraction of the total data available for pre-training.** Line plots showing zero-shot integration performance for each model's embeddings as evaluated by normalized mutual information (NMI), adjusted Rand index (ARI), batch specific average silhouette width (ASW), cell type label ASW, and average BIO (AvgBIO). For each model, the different colors correspond to the downsampling strategy used to generate the data used for pre-training. The dotted line shows the performance of simply using the highly variable genes as an embedding; the dashed line shows the performance of using principal component projections as an embedding; and the brown solid line corresponds to the "non pre-trained" version of each model, where evaluations were done using the randomly initialized weights.

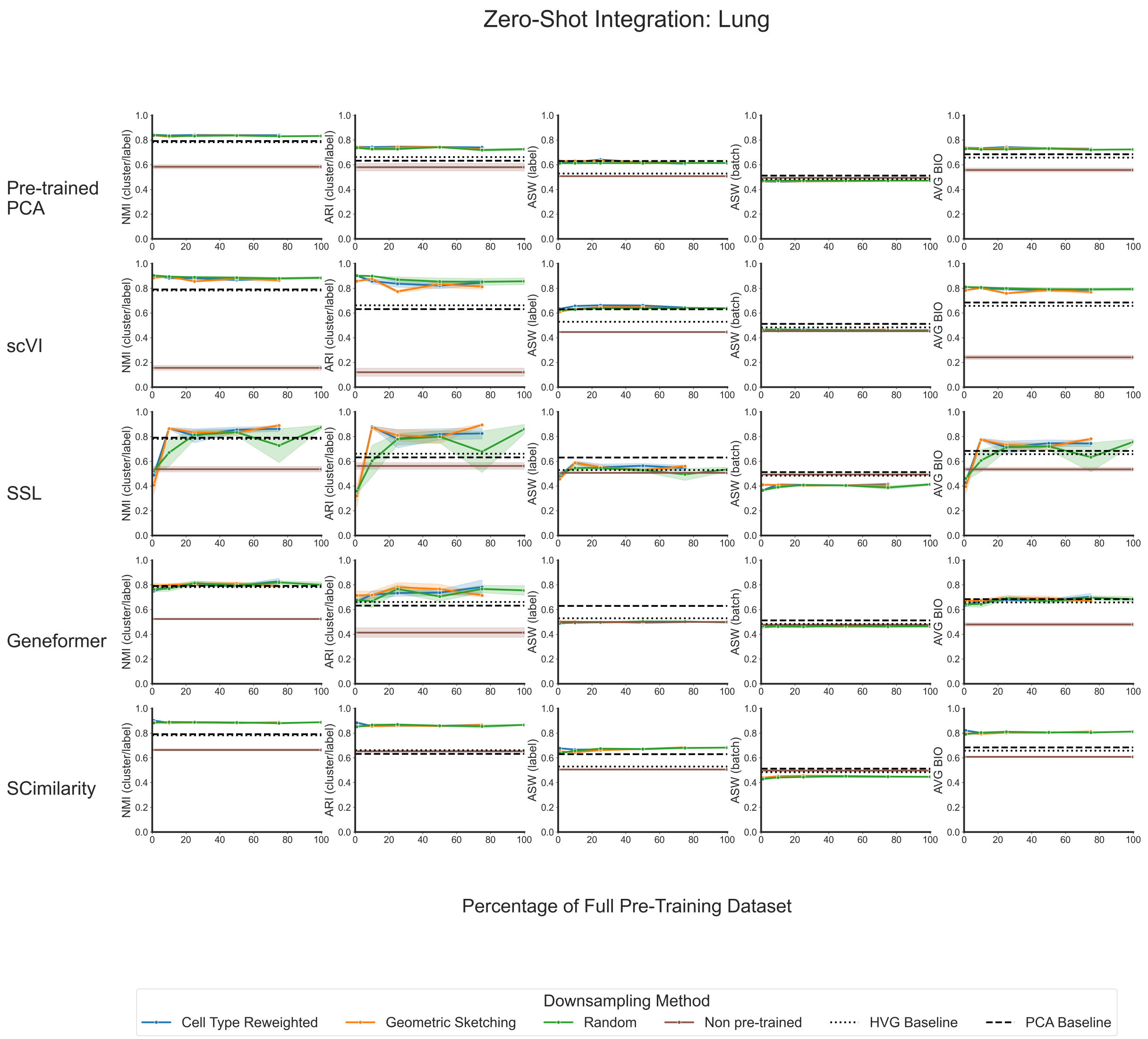


**Supplementary Figure 12. Multiple metrics show that zero-shot model performance on integrating cells from a lung dataset plateaus at a small fraction of the total data available for pre-training.** Line plots showing zero-shot integration performance for each model's embeddings as evaluated by normalized mutual information (NMI), adjusted Rand index (ARI), batch-specific average silhouette width (ASW), cell type label ASW, and average BIO (AvgBIO). For each model, the different colors correspond to the downsampling strategy used to generate the data used for pre-training. The dotted line shows the performance of simply using the highly variable genes as an embedding; the dashed line shows the performance of using principal component projections as an embedding; and the brown solid line corresponds to the "non pre-trained" version of each model, where evaluations were done using the randomly initialized weights.

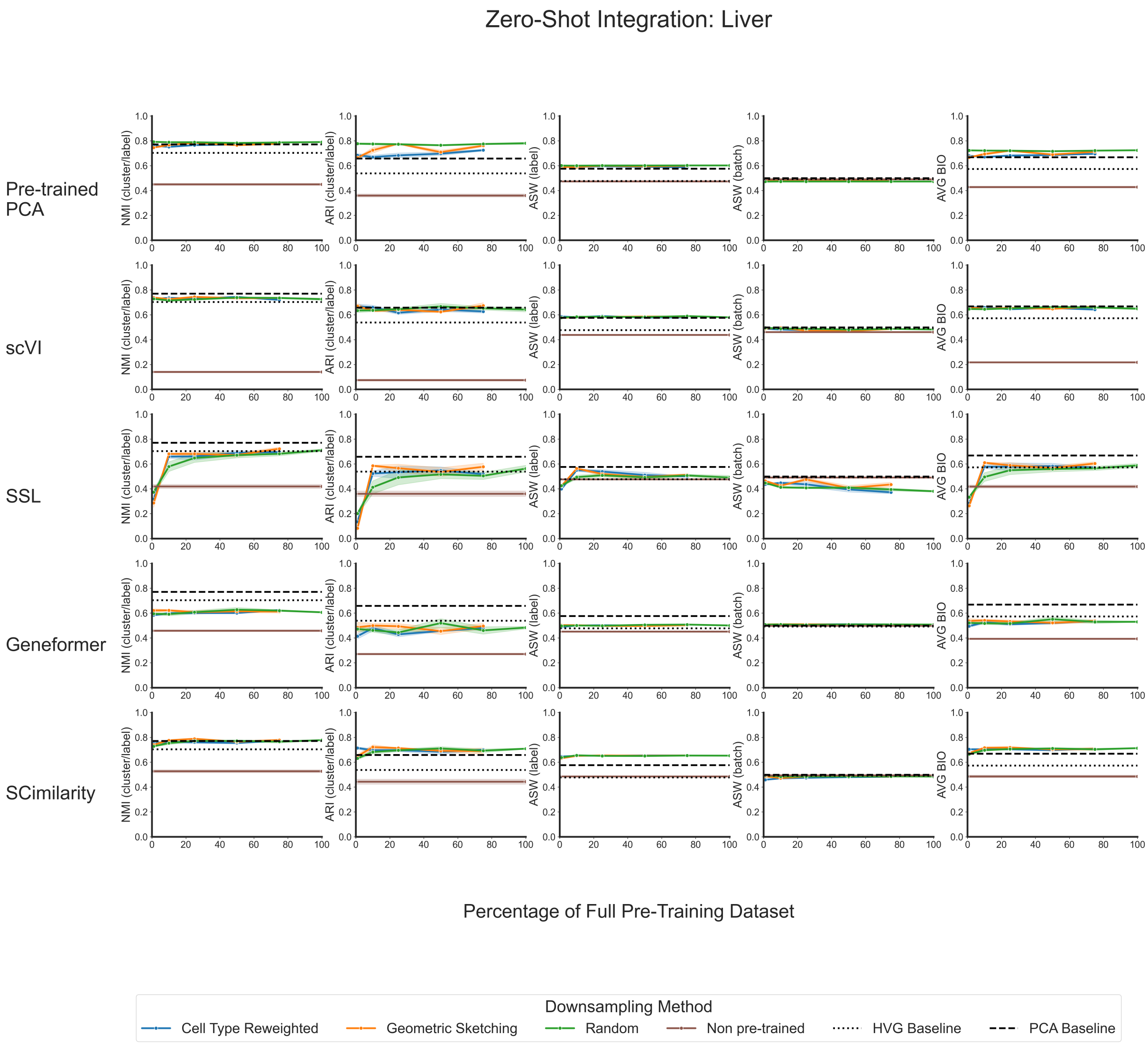


**Supplementary Figure 13. Multiple metrics show that zero-shot model performance on integrating cells from a liver dataset plateaus at a small fraction of the total data available for pre-training.** Line plots showing zero-shot integration performance for each model's embeddings as evaluated by normalized mutual information (NMI), adjusted Rand index (ARI), batch-specific average silhouette width (ASW), cell type label ASW, and average BIO (AvgBIO). For each model, the different colors correspond to the downsampling strategy used to generate the data used for pre-training. The dotted line shows the performance of simply using the highly variable genes as an embedding; the dashed line shows the performance of using principal component projections as an embedding; and the brown solid line corresponds to the "non pre-trained" version of each model, where evaluations were done using the randomly initialized weights.

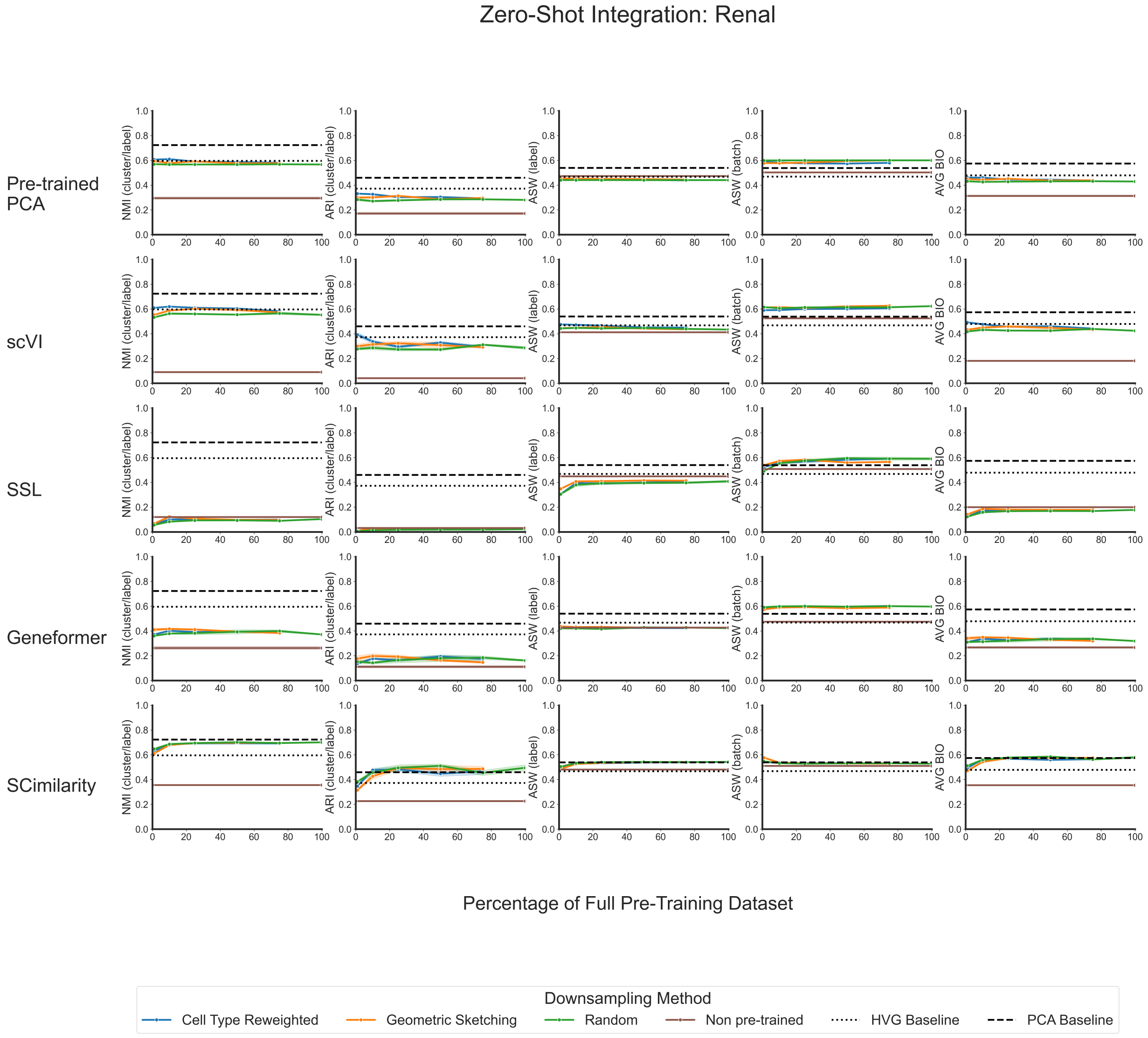


**Supplementary Figure 14. Multiple metrics show that zero-shot model performance on integrating cells from a renal dataset plateaus at a small fraction of the total data available for pre-training.** Line plots showing zero-shot integration performance for each model's embeddings as evaluated by normalized mutual information (NMI), adjusted Rand index (ARI), batch-specific average silhouette width (ASW), cell type label ASW, and average BIO (AvgBIO). For each model, the different colors correspond to the downsampling strategy used to generate the data used for pre-training. The dotted line shows the performance of simply using the highly variable genes as an embedding; the dashed line shows the performance of using principal component projections as an embedding; and the brown solid line corresponds to the "non pre-trained" version of each model, where evaluations were done using the randomly initialized weights.

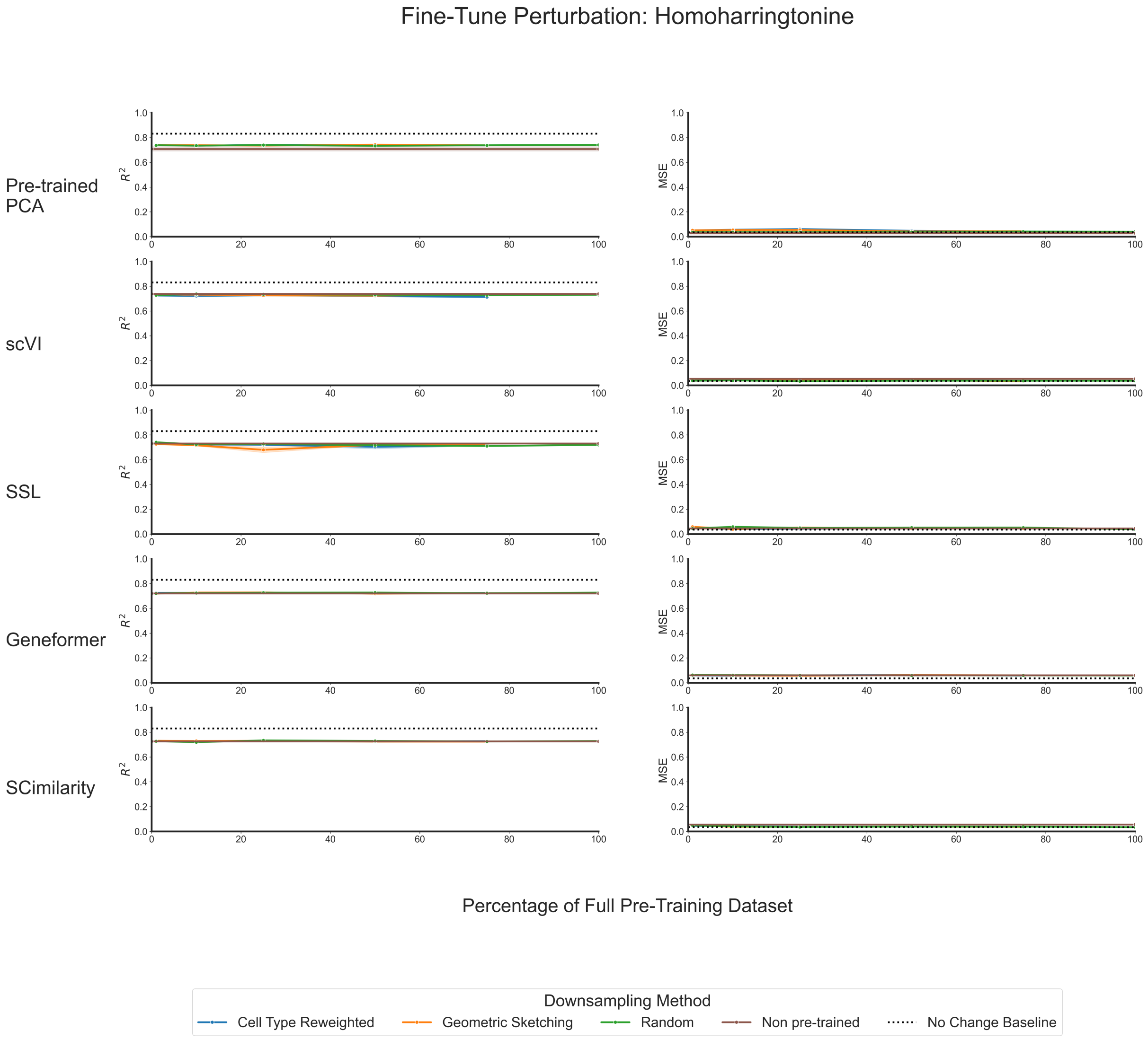


**Supplementary Figure 15. Multiple metrics show that fine-tuned model performance on predicting the gene expression of Homoharringtonine treated cells in the Tahoe-100M dataset plateaus at a small fraction of the total data available for pre-training.** The left hand column depicts line plots showing fine-tuned perturbation performance for each model's embeddings on an out-of-distribution cell line as evaluated by R-squared ($R^2$) between (1) the average predicted gene expression in a perturbed cell and (2) the average observed gene expression in a perturbed cell. The mean squared error (MSE) for the same comparison is given via the line plots in the right hand column. For each model, the different colors correspond to the downsampling strategy used to generate the data used for pre-training. The dotted line shows the performance of simply using the highly variable genes as an embedding; the dashed line shows the performance of using principal component projections as an embedding; and the brown solid line corresponds to the "non pre-trained" version of each model, where evaluations were done using the randomly initialized weights.

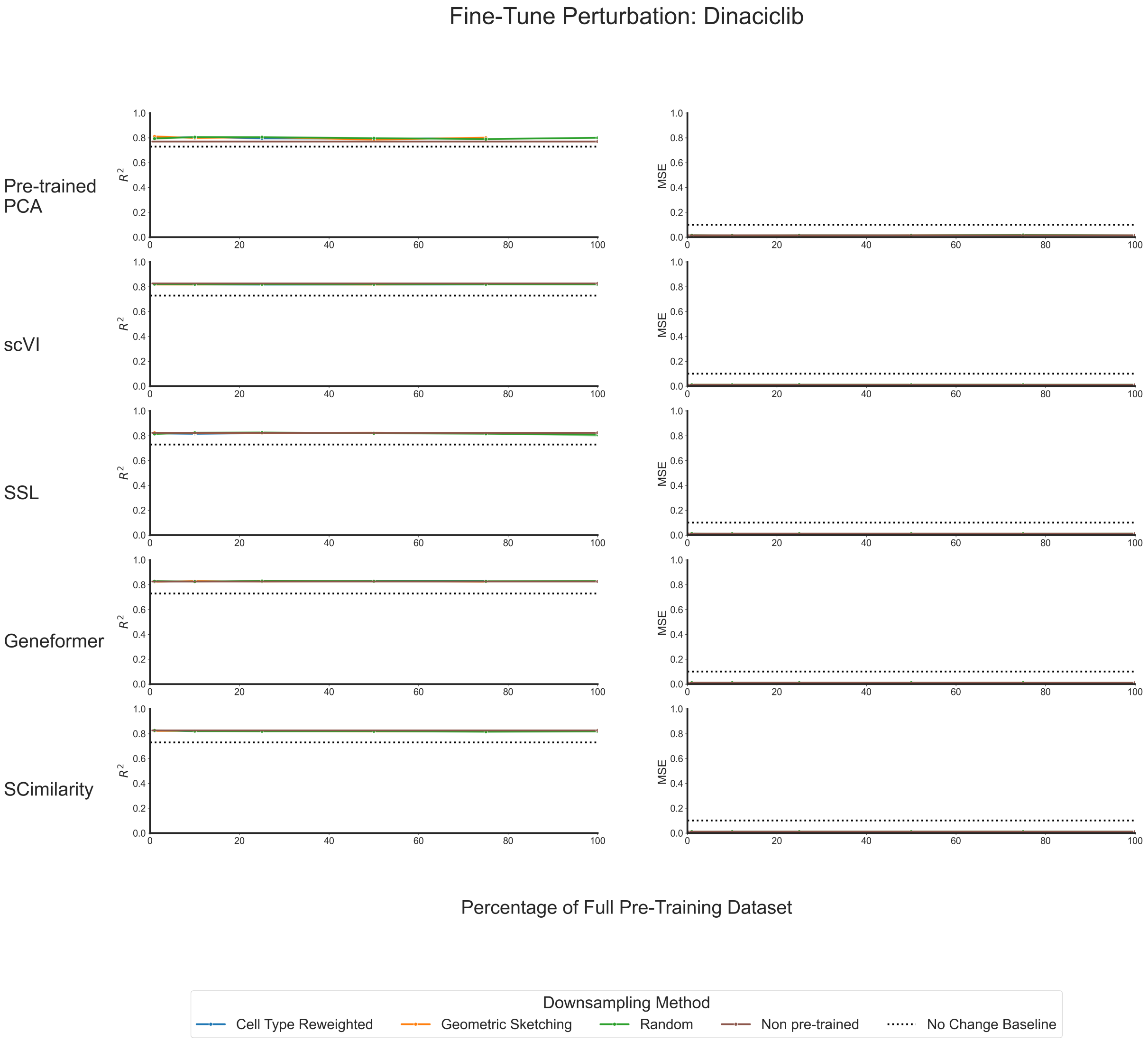


**Supplementary Figure 16. Multiple metrics show that fine-tuned model performance on predicting the gene expression of Dinaciclib treated cells in the Tahoe-100M dataset plateaus at a small fraction of the total data available for pre-training.** The left hand column depicts line plots showing fine-tuned perturbation performance for each model's embeddings on an out-of-distribution cell line as evaluated by R-squared ($R^2$) between (1) the average predicted gene expression in a perturbed cell and (2) the average observed gene expression in a perturbed cell. The mean squared error (MSE) for the same comparison is given via the line plots in the right hand column. For each model, the different colors correspond to the downsampling strategy used to generate the data used for pre-training. The dotted line shows the performance of simply using the highly variable genes as an embedding; the dashed line shows the performance of using principal component projections as an embedding; and the brown solid line corresponds to the "non pre-trained" version of each model, where evaluations were done using the randomly initialized weights.

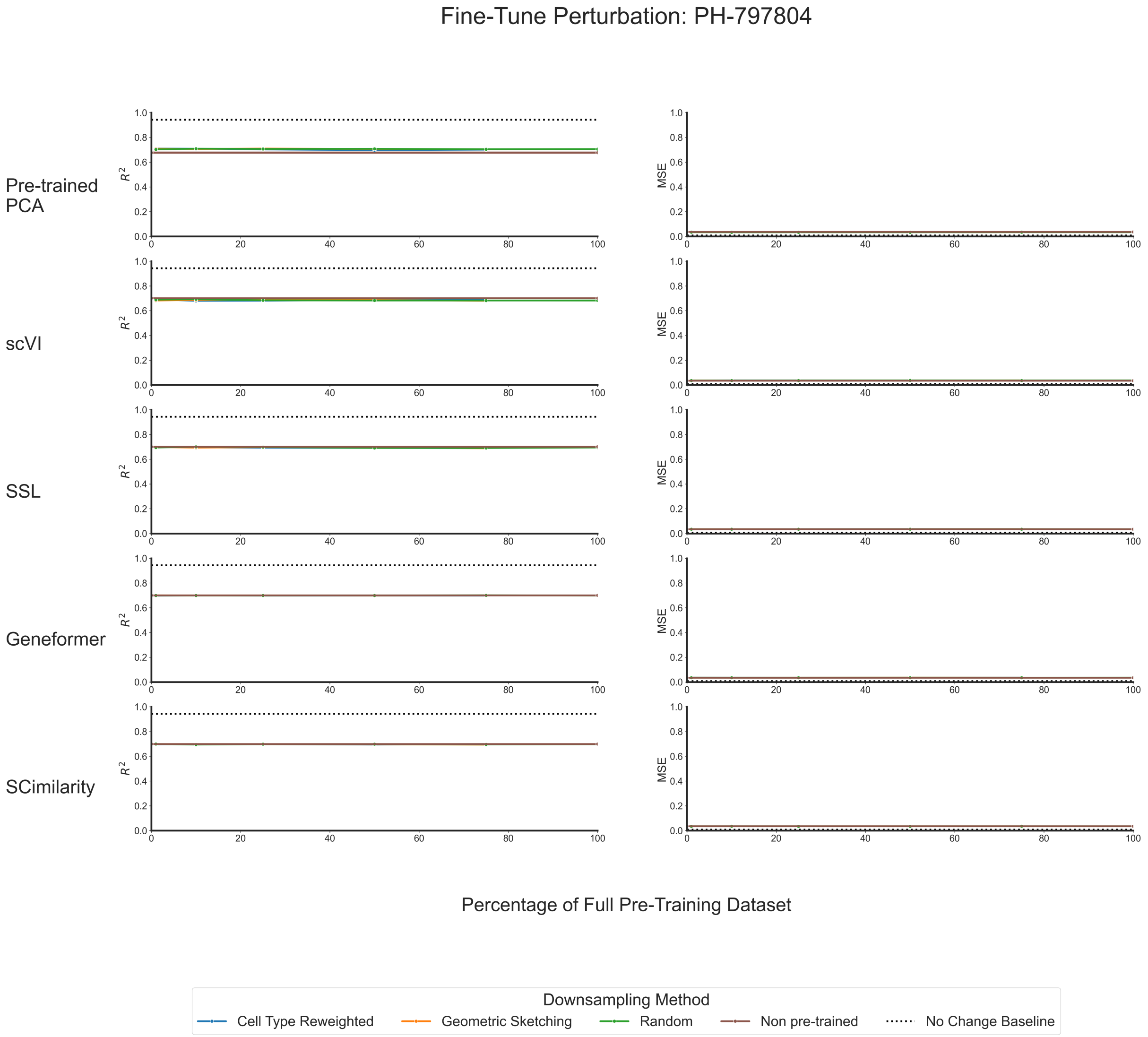


**Supplementary Figure 17. Multiple metrics show that fine-tuned model performance on predicting the gene expression of PH-797804 treated cells in the Tahoe-100M dataset plateaus at a small fraction of the total data available for pre-training.** The left hand column depicts line plots showing fine-tuned perturbation performance for each model's embeddings on an out-of-distribution cell line as evaluated by R-squared ($R^2$) between (1) the average predicted gene expression in a perturbed cell and (2) the average observed gene expression in a perturbed cell. The mean squared error (MSE) for the same comparison is given via the line plots in the right hand column. For each model, the different colors correspond to the downsampling strategy used to generate the data used for pre-training. The dotted line shows the performance of simply using the highly variable genes as an embedding; the dashed line shows the performance of using principal component projections as an embedding; and the brown solid line corresponds to the "non pre-trained" version of each model, where evaluations were done using the randomly initialized weights.

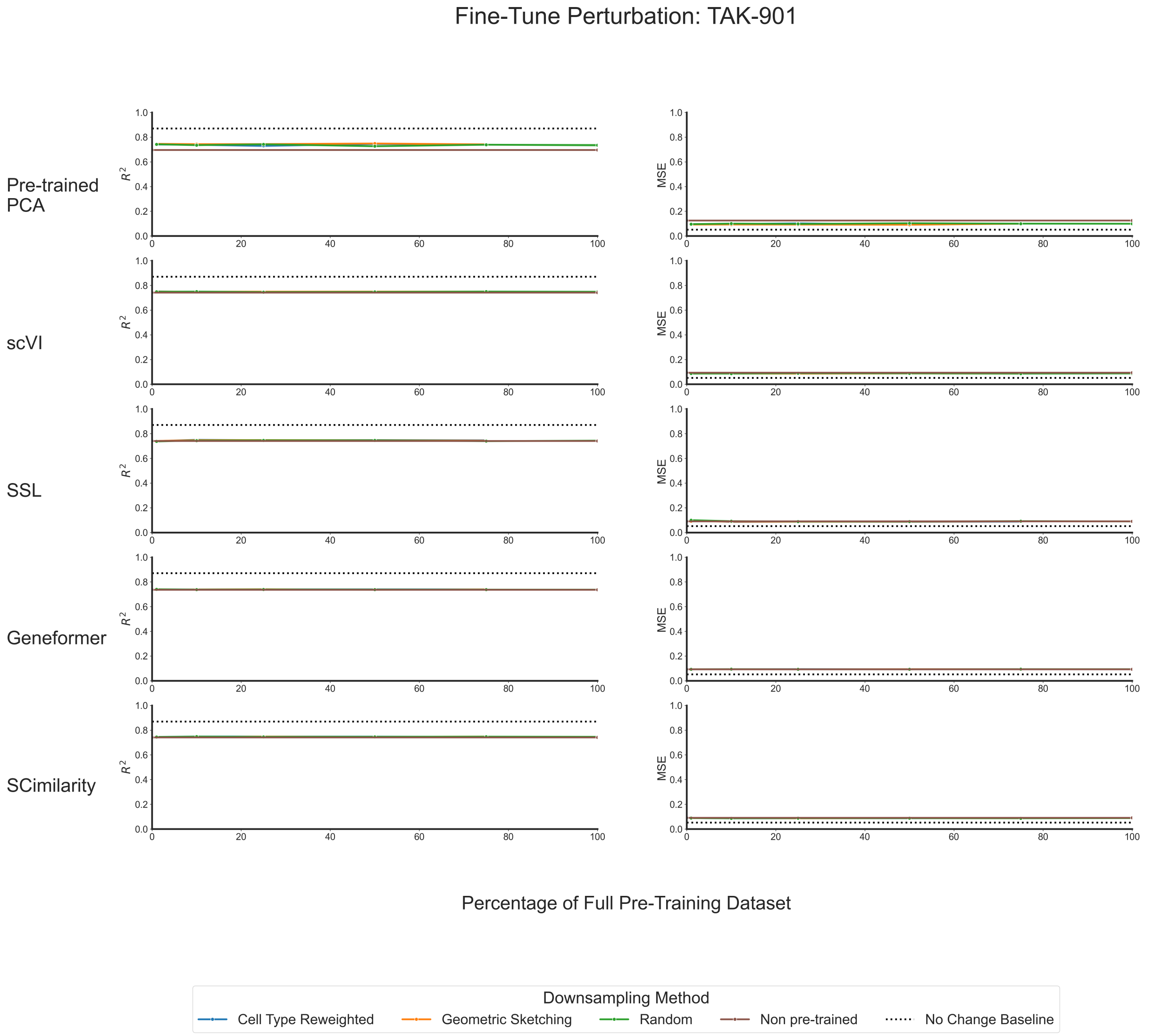


**Supplementary Figure 18. Multiple metrics show that fine-tuned model performance on predicting the gene expression of TAK-901 treated cells in the Tahoe-100M dataset plateaus at a small fraction of the total data available for pre-training.** The left hand column depicts line plots showing fine-tuned perturbation performance for each model's embeddings on an out-of-distribution cell line as evaluated by R-squared ($R^2$) between (1) the average predicted gene expression in a perturbed cell and (2) the average observed gene expression in a perturbed cell. The mean squared error (MSE) for the same comparison is given via the line plots in the right hand column. For each model, the different colors correspond to the downsampling strategy used to generate the data used for pre-training. The dotted line shows the performance of simply using the highly variable genes as an embedding; the dashed line shows the performance of using principal component projections as an embedding; and the brown solid line corresponds to the "non pre-trained" version of each model, where evaluations were done using the randomly initialized weights.

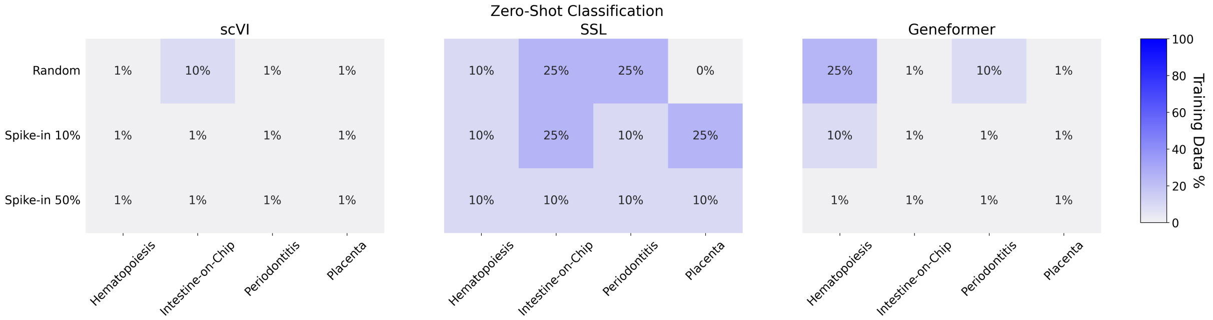


**Supplementary Figure 19. Assessment of zero-shot classification performance saturation as a function of pre-training dataset size across multiple datasets.** A heatmap showing the "learning saturation point" for the clonal hematopoiesis, intestine, periodontitis, and placenta datasets for scVI, SSL, and Geneformer, across each downsampling strategy and when evaluated on classification in the zero-shot case. Each sub-panel corresponds to the model architecture, the $x$-axis corresponds to the dataset evaluated, and the $y$-axis corresponds to the spike-in strategy used to pre-train each model. Thresholds for each model on uniform downsampled data (labeled as "random") are given as reference.

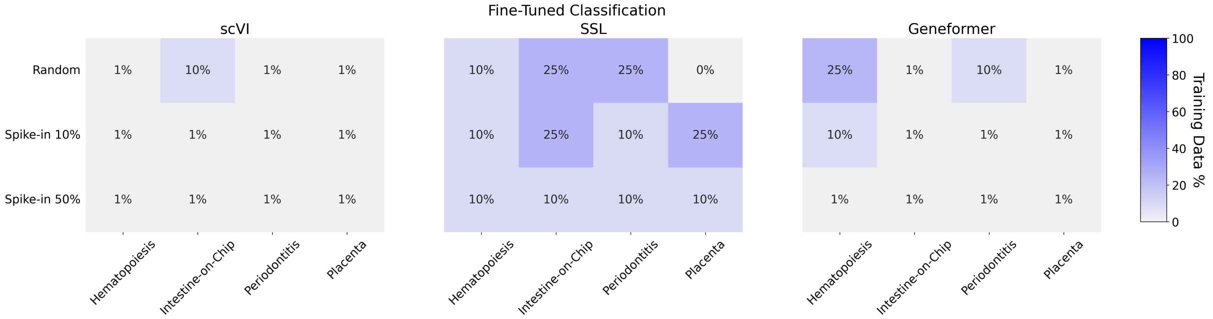


**Supplementary Figure 20. Assessment of fine-tuned classification performance saturation as a function of pre-training dataset size across multiple datasets.** A heatmap showing the "learning saturation point" for the clonal hematopoiesis, intestine, periodontitis, and placenta datasets for scVI, SSL, and Geneformer, across each downsampling strategy and when evaluated on classification in the fine-tuned case. Each sub-panel corresponds to the model architecture, the $x$-axis corresponds to the dataset evaluated, and the $y$-axis corresponds to the spike-in strategy used to pre-train each model. Thresholds for each model on uniform downsampled data (labeled as "random") are given as reference.

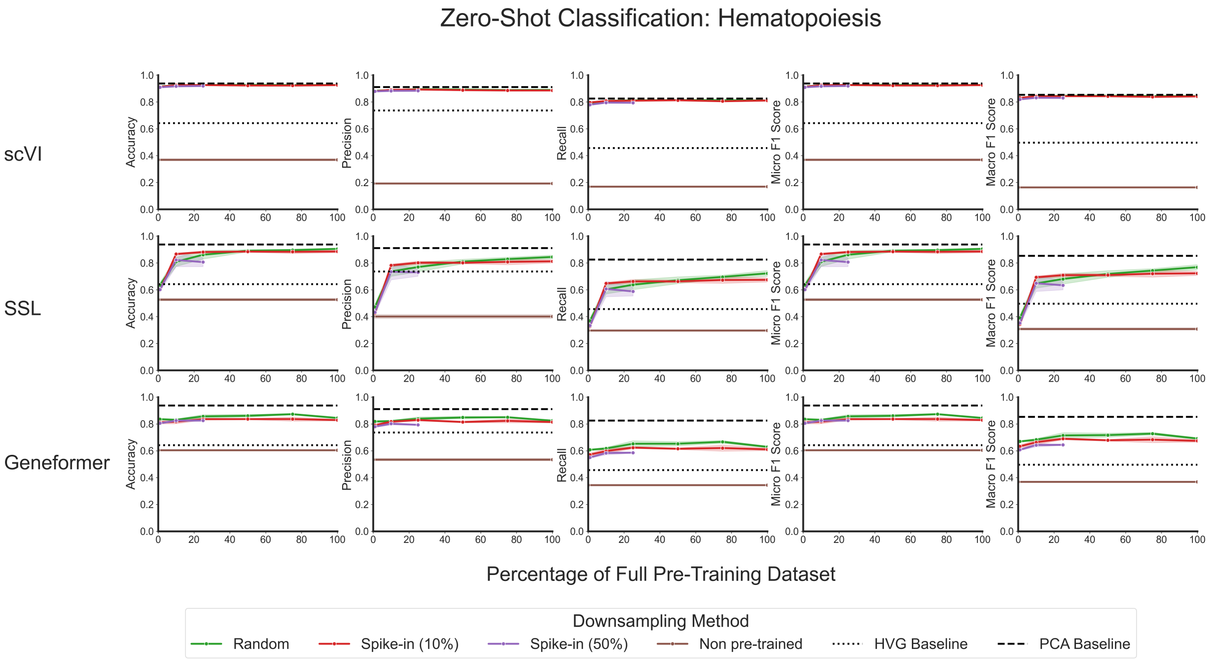


**Supplementary Figure 21. Multiple metrics show that zero-shot model performance on classifying cells from a clonal hematopoiesis dataset plateaus at a small fraction of the total data available for pre-training when perturbation data is spiked in.** Line plots showing zero-shot classification performance for each model's embeddings as evaluated by accuracy, precision, recall, micro F1 score, and macro F1 score. For each model, the different colors correspond to the downsampling strategy used to generate the data used for pre-training. The dotted line shows the performance of simply using the highly variable genes as an embedding; the dashed line shows the performance of using principal component projections as an embedding; and the brown solid line corresponds to the "non pre-trained" version of each model, where evaluations were done using the randomly initialized weights.

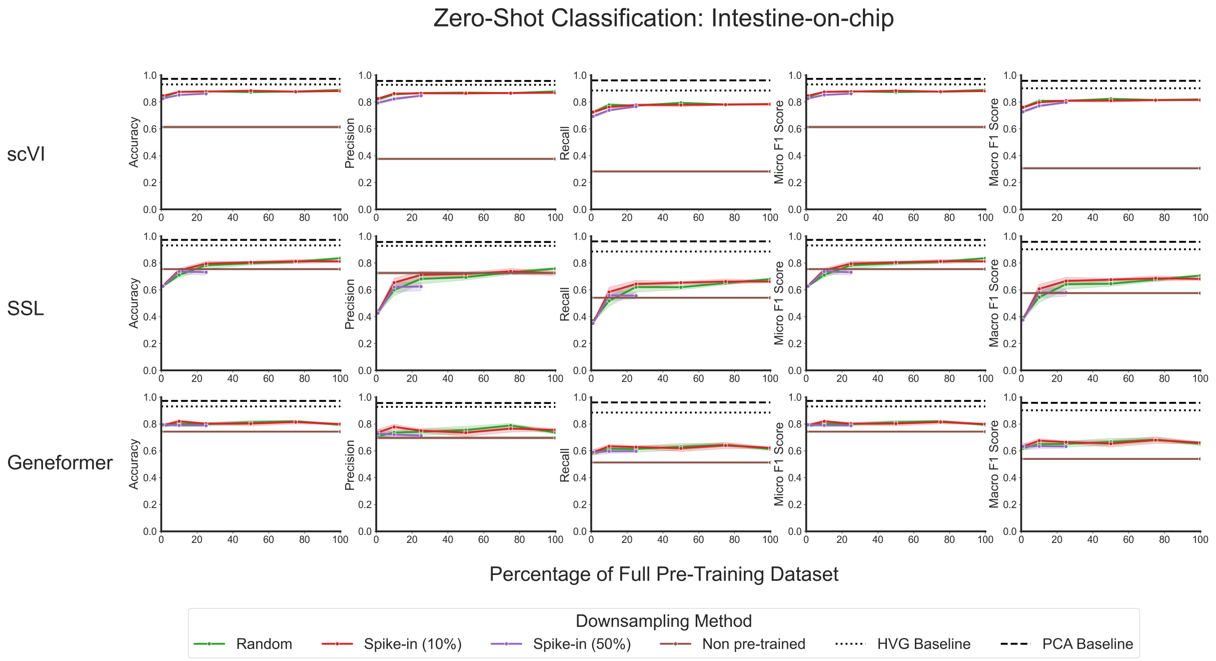


**Supplementary Figure 22. Multiple metrics show that zero-shot model performance on classifying cells from an intestine-on-chip dataset plateaus at a small fraction of the total data available for pre-training when perturbation data is spiked in.** Line plots showing zero-shot classification performance for each model's embeddings as evaluated by accuracy, precision, recall, micro F1 score, and macro F1 score. For each model, the different colors correspond to the downsampling strategy used to generate the data used for pre-training. The dotted line shows the performance of simply using the highly variable genes as an embedding; the dashed line shows the performance of using principal component projections as an embedding; and the brown solid line corresponds to the "non pre-trained" version of each model, where evaluations were done using the randomly initialized weights.

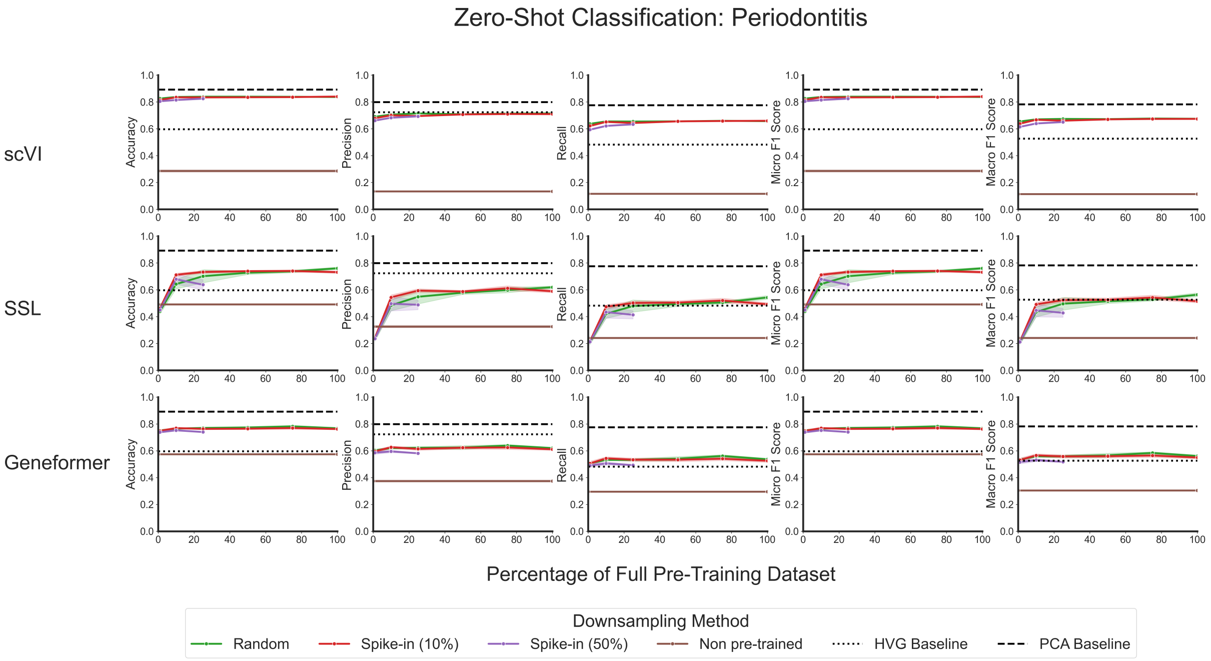


**Supplementary Figure 23. Multiple metrics show that zero-shot model performance on classifying cells from a periodontitis dataset plateaus at a small fraction of the total data available for pre-training when perturbation data is spiked in.** Line plots showing zero-shot classification performance for each model's embeddings as evaluated by accuracy, precision, recall, micro F1 score, and macro F1 score. For each model, the different colors correspond to the downsampling strategy used to generate the data used for pre-training. The dotted line shows the performance of simply using the highly variable genes as an embedding; the dashed line shows the performance of using principal component projections as an embedding; and the brown solid line corresponds to the "non pre-trained" version of each model, where evaluations were done using the randomly initialized weights.

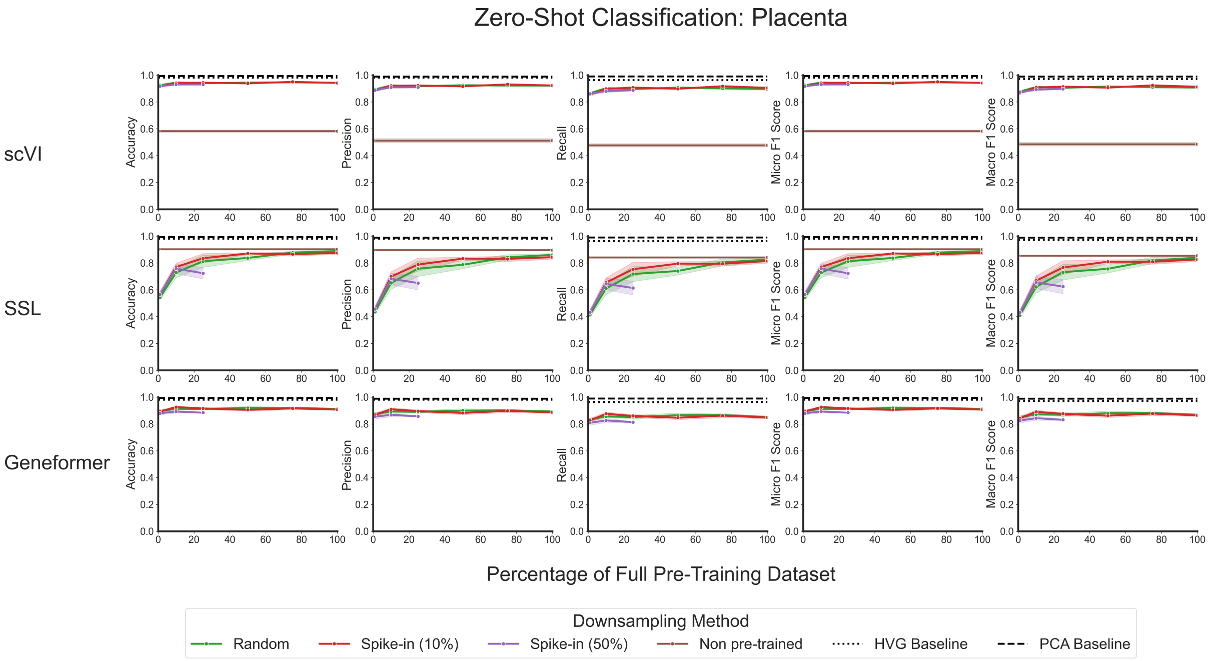


**Supplementary Figure 24. Multiple metrics show that zero-shot model performance on classifying cells from a placenta dataset plateaus at a small fraction of the total data available for pre-training when perturbation data is spiked in.** Line plots showing zero-shot classification performance for each model's embeddings as evaluated by accuracy, precision, recall, micro F1 score, and macro F1 score. For each model, the different colors correspond to the downsampling strategy used to generate the data used for pre-training. The dotted line shows the performance of simply using the highly variable genes as an embedding; the dashed line shows the performance of using principal component projections as an embedding; and the brown solid line corresponds to the "non pre-trained" version of each model, where evaluations were done using the randomly initialized weights.

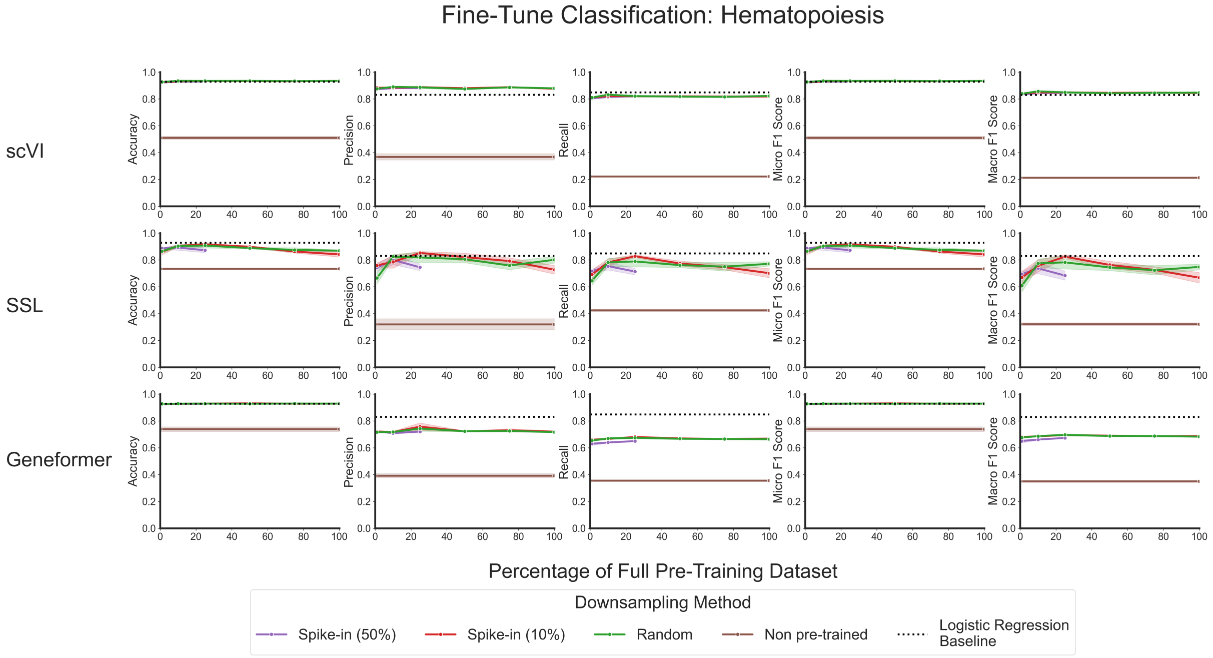


**Supplementary Figure 25. Multiple metrics show that fine-tuned model performance on classifying cells from a clonal hematopoiesis dataset plateaus at a small fraction of the total data available for pre-training when perturbation data is spiked in.** Line plots showing zero-shot classification performance for each model's embeddings as evaluated by accuracy, precision, recall, micro F1 score, and macro F1 score. For each model, the different colors correspond to the downsampling strategy used to generate the data used for pre-training. The dotted line shows the performance of a regularized logistic classifier using the highly variable genes as input; and the brown solid line corresponds to the "non pre-trained" version of each model, where evaluations were done using the randomly initialized weights.

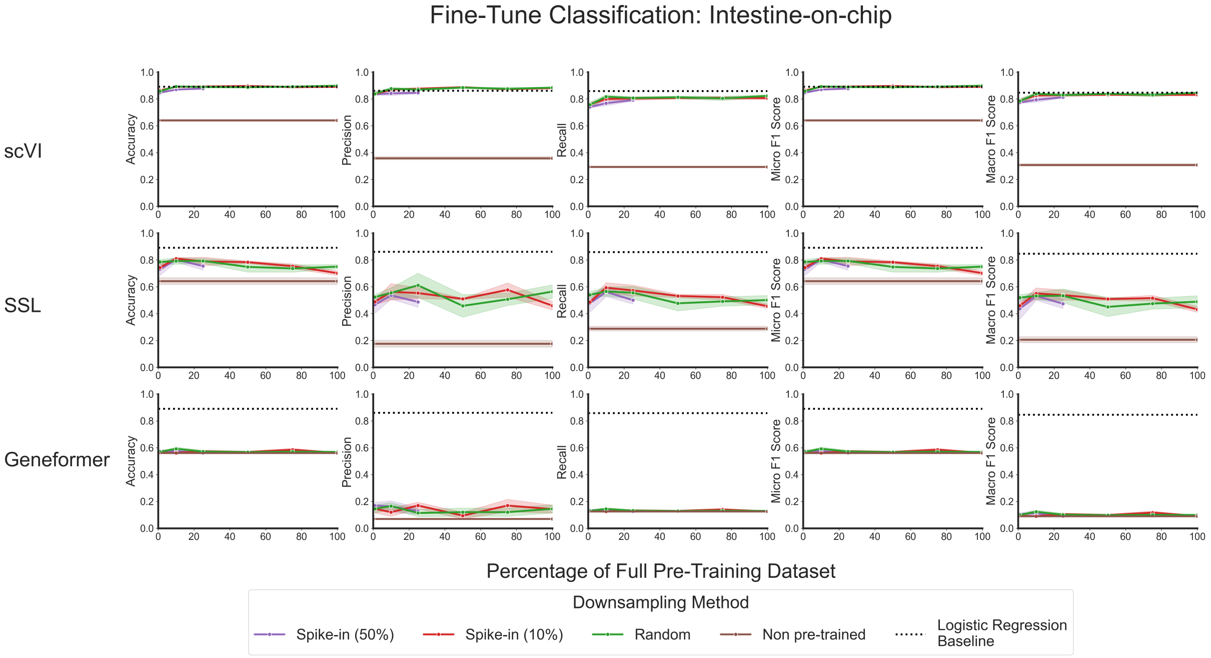


**Supplementary Figure 26. Multiple metrics show that fine-tuned model performance on classifying cells from an intestine-on-chip dataset plateaus at a small fraction of the total data available for pre-training when perturbation data is spiked in.** Line plots showing zero-shot classification performance for each model's embeddings as evaluated by accuracy, precision, recall, micro F1 score, and macro F1 score. For each model, the different colors correspond to the downsampling strategy used to generate the data used for pre-training. The dotted line shows the performance of a regularized logistic classifier using the highly variable genes as input; and the brown solid line corresponds to the "non pre-trained" version of each model, where evaluations were done using the randomly initialized weights.

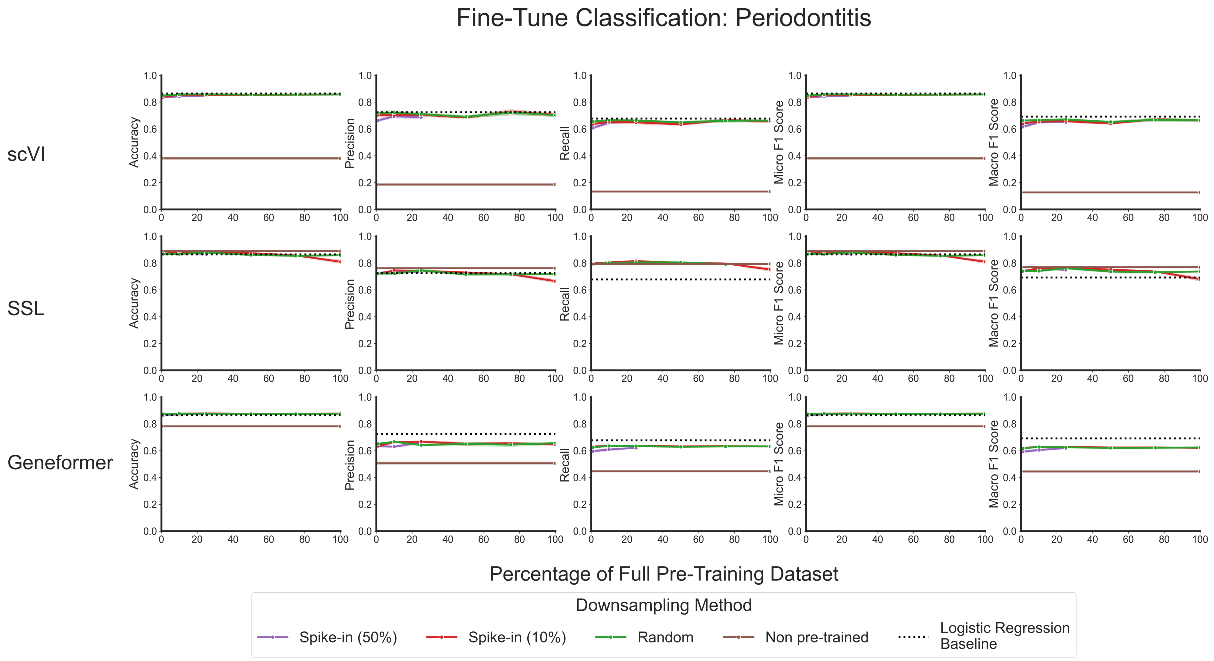


**Supplementary Figure 27. Multiple metrics show that fine-tuned model performance on classifying cells from a periodontitis dataset plateaus at a small fraction of the total data available for pre-training when perturbation data is spiked in.** Line plots showing zero-shot classification performance for each model's embeddings as evaluated by accuracy, precision, recall, micro F1 score, and macro F1 score. For each model, the different colors correspond to the downsampling strategy used to generate the data used for pre-training. The dotted line shows the performance of a regularized logistic classifier using the highly variable genes as input; and the brown solid line corresponds to the "non pre-trained" version of each model, where evaluations were done using the randomly initialized weights.

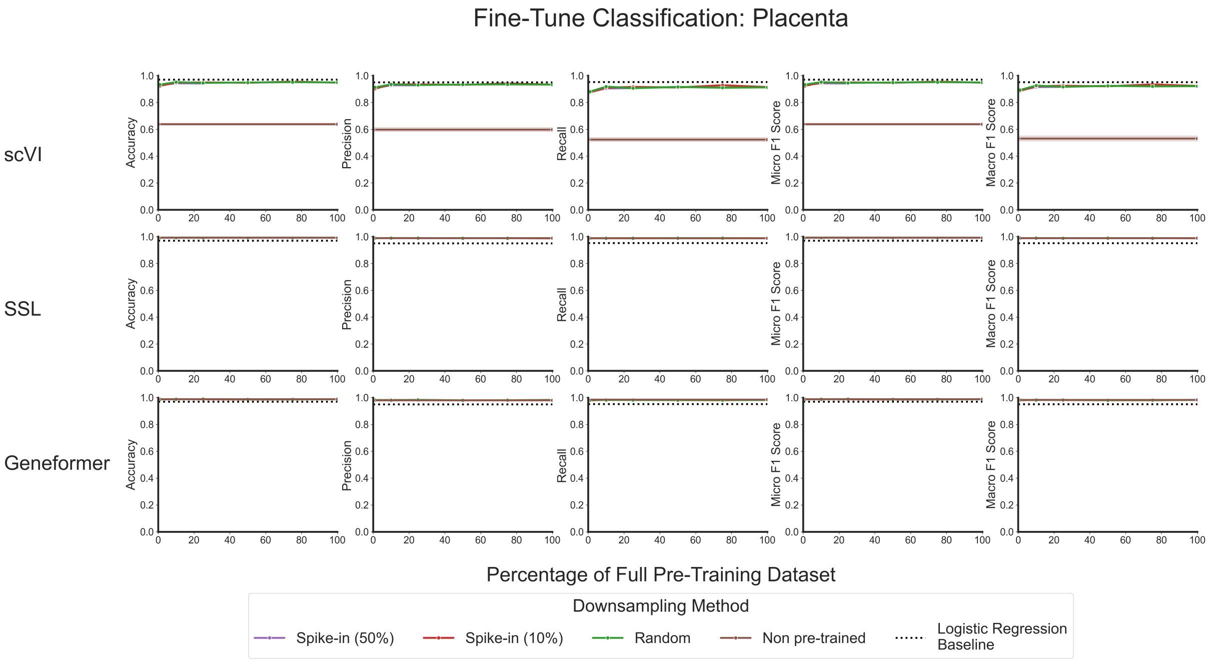


**Supplementary Figure 28. Multiple metrics show that fine-tuned model performance on classifying cells from a placenta dataset plateaus at a small fraction of the total data available for pre-training when perturbation data is spiked in.** Line plots showing zero-shot classification performance for each model's embeddings as evaluated by accuracy, precision, recall, micro F1 score, and macro F1 score. For each model, the different colors correspond to the downsampling strategy used to generate the data used for pre-training. The dotted line shows the performance of a regularized logistic classifier using the highly variable genes as input; and the brown solid line corresponds to the "non pre-trained" version of each model, where evaluations were done using the randomly initialized weights.

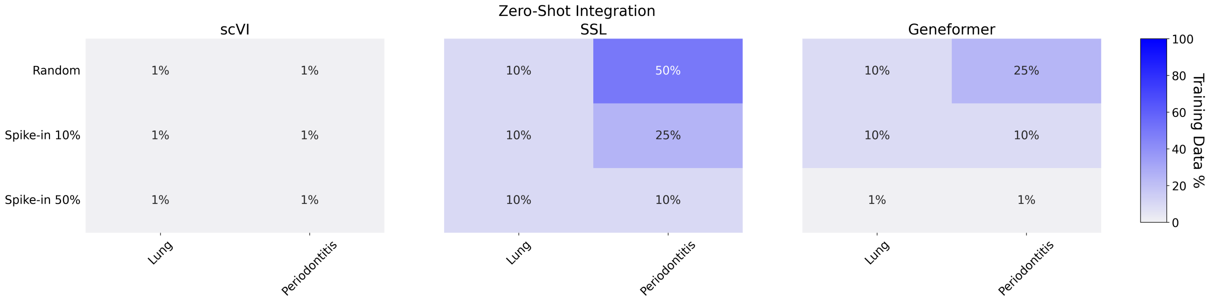


**Supplementary Figure 29. Assessment of zero-shot integration performance saturation as a function of pre-training dataset size across multiple datasets.** A heatmap showing the "learning saturation point" for the periodontitis dataset and the lung dataset for scVI, SSL, and Geneformer, across each downsampling strategy and when evaluated on zero-shot batch integration case. Each sub-panel corresponds to the model architecture, the $x$-axis corresponds to the dataset evaluated, and the $y$-axis corresponds to the spike-in strategy used to pre-train each model. Thresholds for each model on uniform downsampled data (labeled as "random") are given as reference.

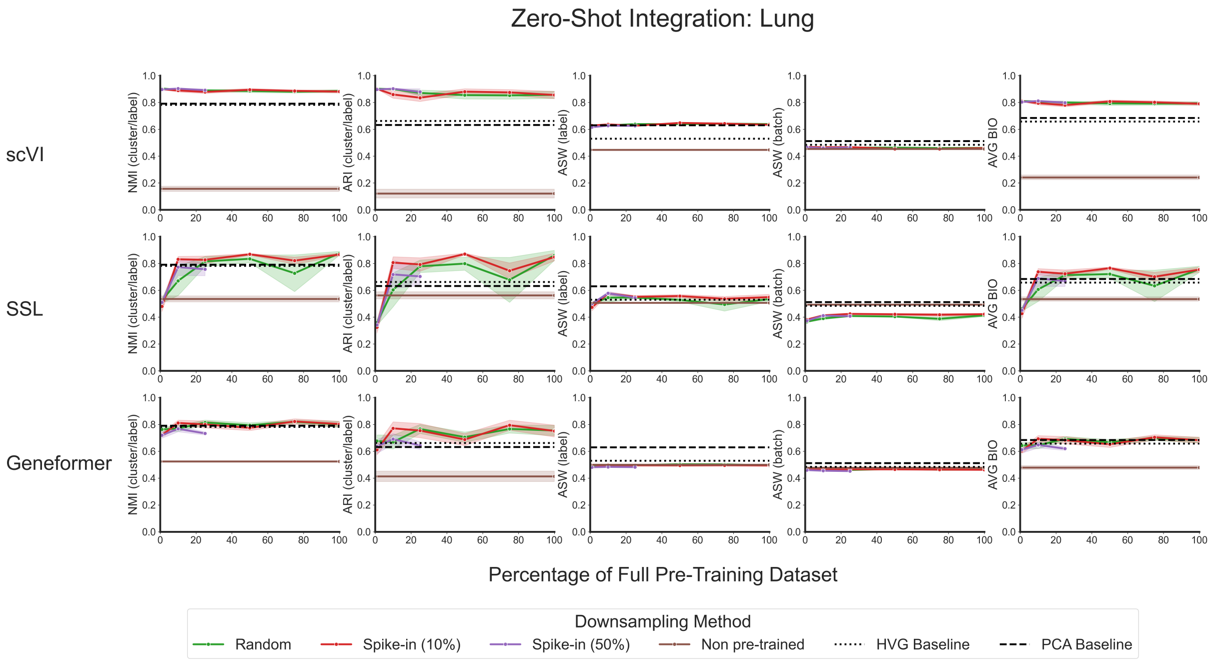


**Supplementary Figure 30. Multiple metrics show that zero-shot model performance on integrating cells from a lung dataset plateaus at a small fraction of the total data available for pre-training when perturbation data is spiked in.** Line plots showing zero-shot integration performance for each model's embeddings as evaluated by normalized mutual information (NMI), adjusted Rand index (ARI), batch-specific average silhouette width (ASW), cell type label ASW, and average BIO (AvgBIO). For each model, the different colors correspond to the downsampling strategy used to generate the data used for pre-training. The dotted line shows the performance of simply using the highly variable genes as an embedding; the dashed line shows the performance of using principal component projections as an embedding; and the brown solid line corresponds to the "non pre-trained" version of each model, where evaluations were done using the randomly initialized weights.

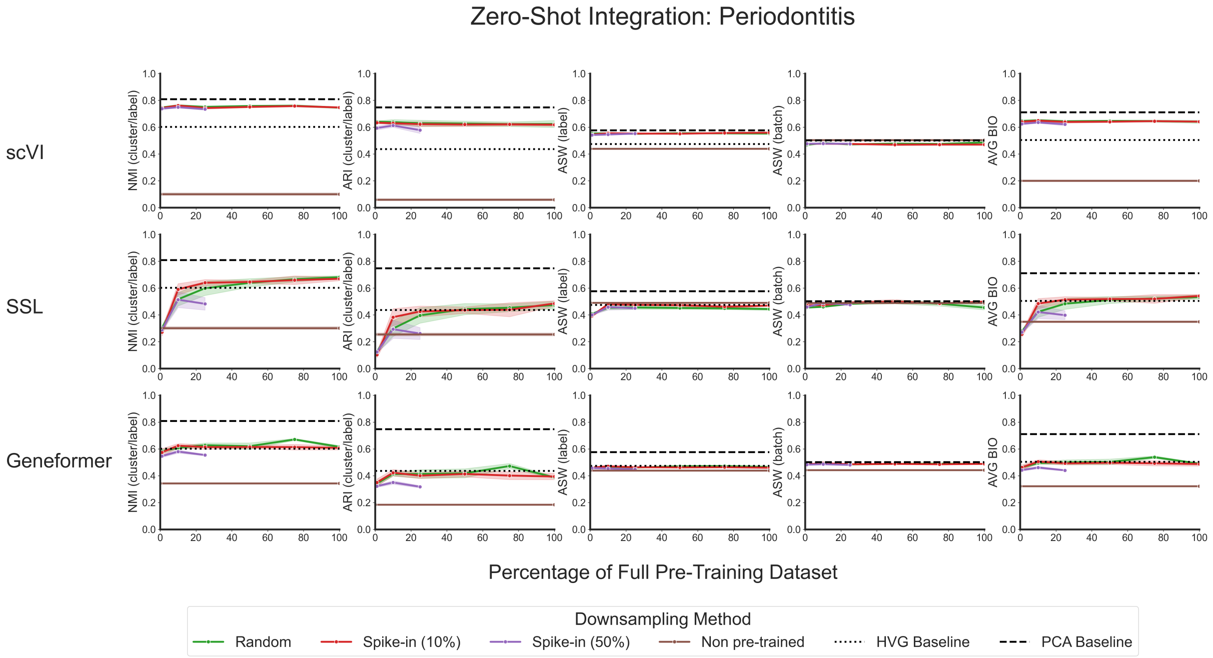


**Supplementary Figure 31. Multiple metrics show that zero-shot model performance on integrating cells from a periodontitis dataset plateaus at a small fraction of the total data available for pre-training when perturbation data is spiked in.** Line plots showing zero-shot integration performance for each model's embeddings as evaluated by normalized mutual information (NMI), adjusted Rand index (ARI), batch-specific average silhouette width (ASW), cell type label ASW, and average BIO (AvgBIO). For each model, the different colors correspond to the downsampling strategy used to generate the data used for pre-training. The dotted line shows the performance of simply using the highly variable genes as an embedding; the dashed line shows the performance of using principal component projections as an embedding; and the brown solid line corresponds to the "non pre-trained" version of each model, where evaluations were done using the randomly initialized weights.

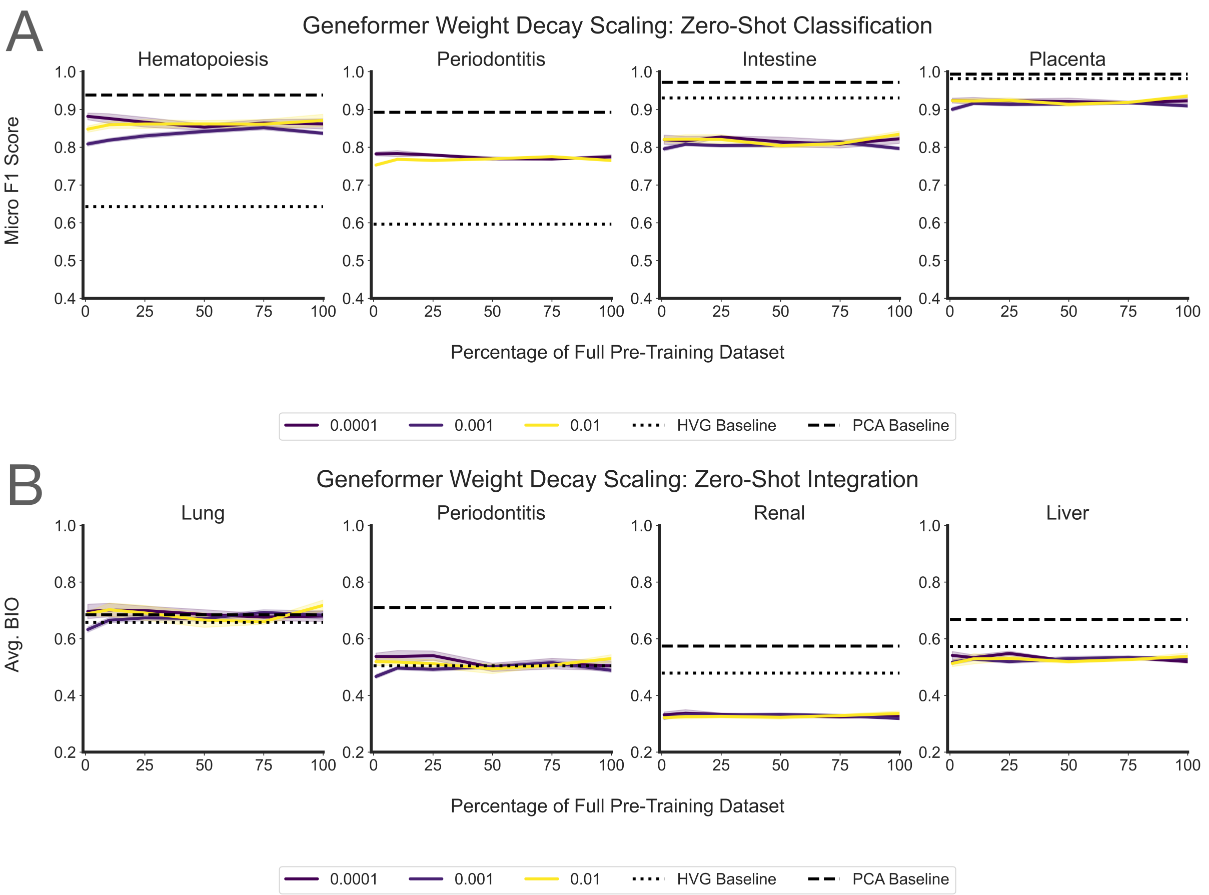


**Supplementary Figure 32. Evaluation of the impact of varying hyperparameters when training Geneformer.** Line plots showing zero-shot classification **(A)** and zero-shot integration **(B)** performance for each Geneformer model's embeddings. Zero-shot classification is evaluated by accuracy, precision, recall, micro F1 score, and macro F1 score. Zero-shot integration is evaluated by normalized mutual information (NMI), adjusted Rand index (ARI), batch-specific average silhouette width (ASW), cell type label ASW, and average BIO (AvgBIO). For each model, the different colors correspond to varying weight decay hyperparameters used during pre-training. The dotted line shows the performance of simply using the highly variable genes as an embedding, and the dashed line shows the performance of using principal component projections as an embedding.

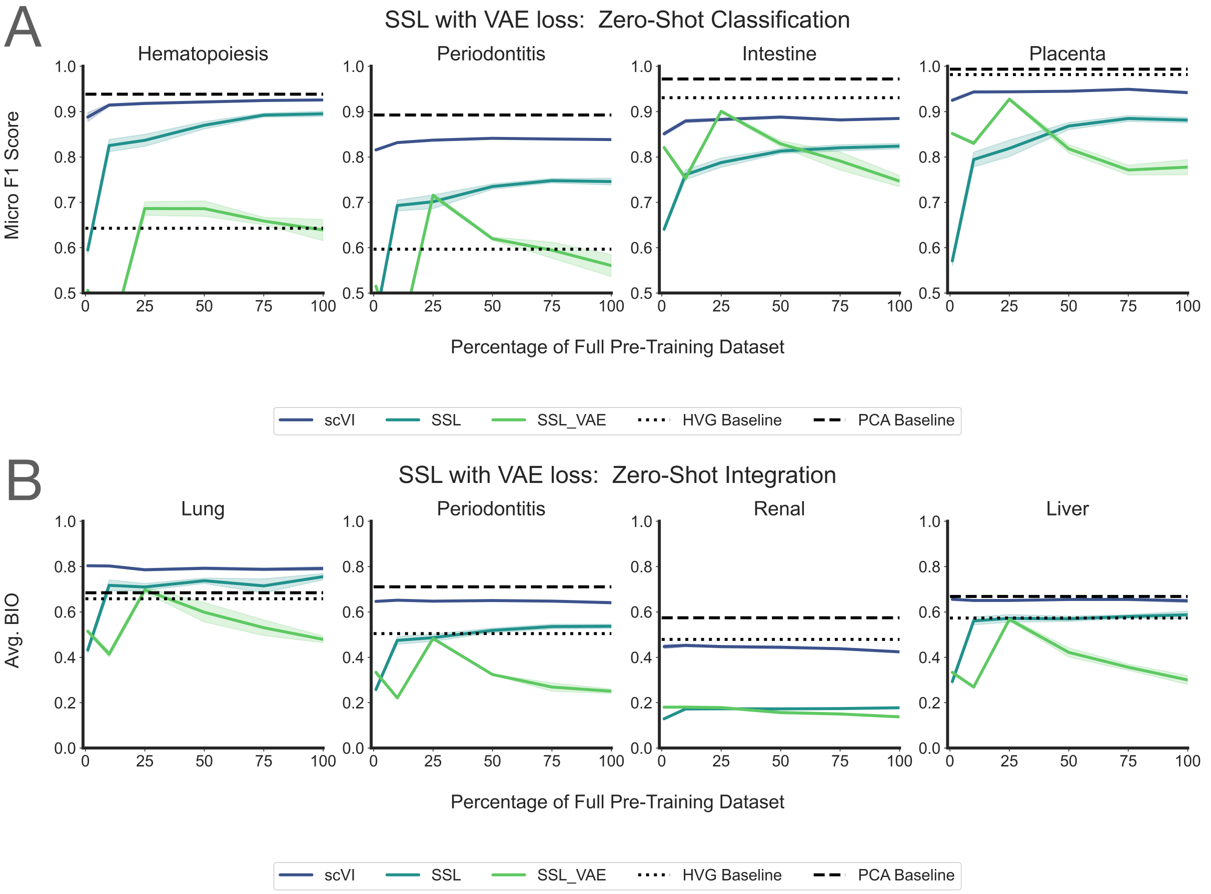


**Supplementary Figure 33. Evaluation of the impact of combining the VAE loss used in scVI with the masked training objective used in SSL.** Line plots showing zero-shot classification **(A)** and zero-shot integration **(B)** performance for each model's embeddings from scVI, SSL, and SSL trained with an additional VAE loss (SSL_VAE). Zero-shot classification is evaluated by accuracy, precision, recall, micro F1 score, and macro F1 score. Zero-shot integration is evaluated by normalized mutual information (NMI), adjusted Rand index (ARI), batch-specific average silhouette width (ASW), cell type label ASW, and average BIO (AvgBIO). For each model, the different colors correspond to varying weight decay hyperparameters used during pre-training. The dotted line shows the performance of simply using the highly variable genes as an embedding, and the dashed line shows the performance of using principal component projections as an embedding.

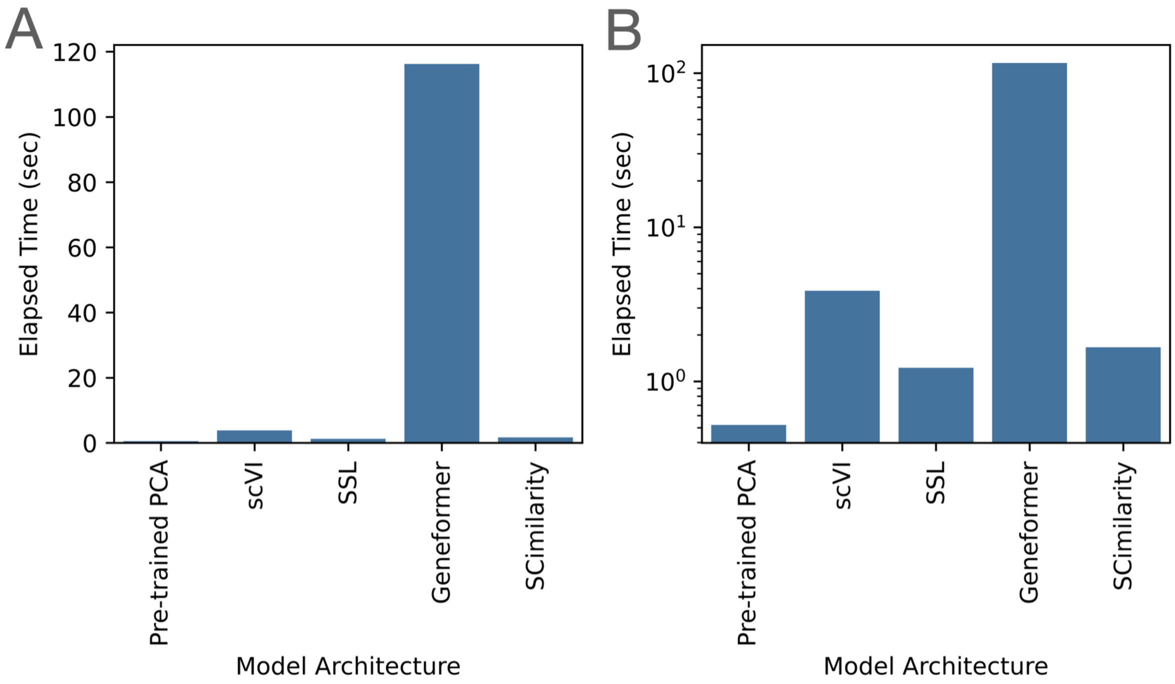


**Supplementary Figure 34. Evaluation of the computational cost of inference for each model architecture.** Each barplot shows the wall clock time needed for Pre-trained PCA, scVI, SSL, and Geneformer to compute model embeddings for 10K cells. The result for Geneformer also includes the time needed to tokenize the input data. **(A)** Time taken (in seconds) for each model architecture. **(B)** The same as **(A)**, but with the time taken plotted on a log scale.
